# Adipocyte lipin 1 is positively associated with metabolic health in humans and regulates systemic metabolism in mice

**DOI:** 10.1101/2023.02.01.526676

**Authors:** Andrew LaPoint, Jason M. Singer, Daniel Ferguson, Trevor M. Shew, M. Katie Renkemeyer, Hector Palacios, Rachael Field, Mahalakshmi Shankaran, Gordon I. Smith, Jun Yoshino, Mai He, Gary J. Patti, Marc K. Hellerstein, Samuel Klein, Jonathan R. Brestoff, Brian N. Finck, Andrew J. Lutkewitte

## Abstract

Dysfunctional adipose tissue is believed to promote the development of hepatic steatosis and systemic insulin resistance, but many of the mechanisms involved are still unclear. Lipin 1 catalyzes the conversion of phosphatidic acid to diacylglycerol (DAG), the penultimate step of triglyceride synthesis, which is essential for lipid storage. Herein we found that adipose tissue *LPIN1* expression is decreased in people with obesity compared to lean subjects and low *LPIN1* expression correlated with multi-tissue insulin resistance and increased rates of hepatic de novo lipogenesis. Comprehensive metabolic and multi-omic phenotyping demonstrated that adipocyte-specific *Lpin1-/-* mice had a metabolically-unhealthy phenotype, including liver and skeletal muscle insulin resistance, hepatic steatosis, increased hepatic de novo lipogenesis, and transcriptomic signatures of nonalcoholic steatohepatitis that was exacerbated by high-fat diets. We conclude that adipocyte lipin 1-mediated lipid storage is vital for preserving adipose tissue and systemic metabolic health and its loss predisposes mice to nonalcoholic steatohepatitis.

## Main

Adipose tissue plays important roles in maintaining metabolic homeostasis by serving as a reservoir to store fat (lipogenesis) and as a secretory organ that releases hormones and other biological factors that signal to distal organs. Although increases in adipose tissue mass (obesity) are associated with an increased risk for hepatic steatosis, systemic insulin resistance, and cardiometabolic diseases, a subset of people with obesity do not develop these abnormalities and have metabolically normal obesity (MHO), as opposed to metabolically unhealthy obesity (MUO)^1–3^. These studies suggest that dysfunctional adipose tissue (defined here by adipose tissue insulin resistance, inflammation, and fibrosis), not adipose tissue mass, is linked to metabolic disease development. For example, whole body insulin resistance is associated with decreased adipose tissue lipogenic gene expression in people with MUO compared to people with metabolically MNO^3–6^. Alternatively, enhanced lipogenic capacity (i.e., lipid storage capacity) in adipose tissue prevents systemic insulin resistance in mouse models of obesity and is associated with increased insulin sensitivity in people^3,7,8^. Furthermore, adipose tissue inflammation and fibrosis are also associated with insulin resistance and hepatic steatosis in people with obesity^9^.

Adipocytes store lipids as triglycerides (TAG) by the sequential acylation of glycerol-3-phosphate. The penultimate step of TAG synthesis, which is the conversion of phosphatidic acid to diacylglycerol (DAG), is carried out by phosphatidic acid phosphohydrolase (PAP) enzymes known as lipins^10,11^. There are three lipin family proteins (lipin 1, 2, 3) that are differentially expressed in various tissues^12,13^. *Lpin1*, which encodes lipin 1, was first identified as the gene deleted in fatty liver dystrophic (*fld*) mice^12^, which exhibit severe lipodystrophy that is associated with systemic metabolic abnormalities^14,15^. *Lpin1* is highly expressed in adipocytes and is essential for TAG synthesis in mice through its action as a PAP enzyme. Lipin 1 is also required for the initiation of the adipogenic gene expression program, likely via the effects of phosphatidic acid on signaling cascades and direct transcriptional regulation of gene expression in the nucleus^16–20^. Conditional mice with adipocyte-specific expression of a truncated form of lipin 1, which lacked intrinsic PAP activity but retained its activity as a transcriptional regulator, exhibited a lean phenotype but were not severely lipodystrophic^21,22^. These mice also exhibited signs of insulin resistance despite being lean^21,22^, whereas lipin 1-overexpressing mice had increased adiposity, yet were insulin sensitive^23^. Prior work has also identified correlations between high adipose tissue *LPIN1* expression and insulin sensitivity in people^24–26^. Collectively, these studies suggest that high expression of lipin 1 in adipocytes may preserve adipose tissue function and protect mice from the development of fatty liver, insulin resistance, and other metabolic abnormalities.

Despite these striking effects of modulating adipocyte lipin 1 activity on metabolic health, the mechanisms by which lipin 1 regulates systemic metabolism, specifically through adipose tissue-liver cross-talk are poorly understood. Moreover, prior studies of lipin 1 loss of function models were complicated by the expression of the truncated lipin 1 protein that retained activity as a transcriptional regulator^21^. Herein, we used mice with a floxed *Lpin1* allele that was designed to completely delete lipin 1 protein^27,28^ to generate mice with total loss of lipin 1 in adipocytes and characterized their metabolic phenotype by using rigorous metabolic phenotyping and multiomic approaches. In addition, we evaluated the expression of *LPIN1* in subcutaneous adipose tissue obtained from people who were healthy and lean and those with MHO and MUO.

## Results

### Adipose tissue *LPIN1* expression is decreased in people with metabolically unhealthy obesity and correlates with insulin sensitivity and hepatic DNL

We compared the expression of *LPIN1* in subcutaneous abdominal adipose tissue (SAAT) from people who are metabolically healthy and lean (MHL), metabolically healthy with obesity (MHO), and metabolically unhealthy with obesity (MUO), stratified based on body mass index, glucose tolerance, HbA1c and IHTG content assessed by MRI (Data Table 1). Expression of *LPIN1* was highest in the MHL group, and progressively decreased from MHL to the MHO to the MUO group (Fig. 1a). In addition, *LPIN1* expression directly correlated with skeletal muscle insulin sensitivity, defined as the glucose disposal rate [Rd] per kg fat-free mass (FFM) divided by plasma insulin during a hyperinsulinemic-euglycemic clamp procedure^29^ and hepatic insulin sensitivity (assessed as the Hepatic Insulin Sensitivity Index (HISI), which is the reciprocal of the product of basal endogenous glucose production rate and basal plasma insulin concentration)^30,31^. In contrast, *LPIN1* expression was inversely associated with rates of hepatic *de novo* lipogenesis (DNL) (Fig.1b-d). These data demonstrate that adipose tissue *LPIN1* expression is associated with a healthy metabolic profile in people.

**Data Table 1.**
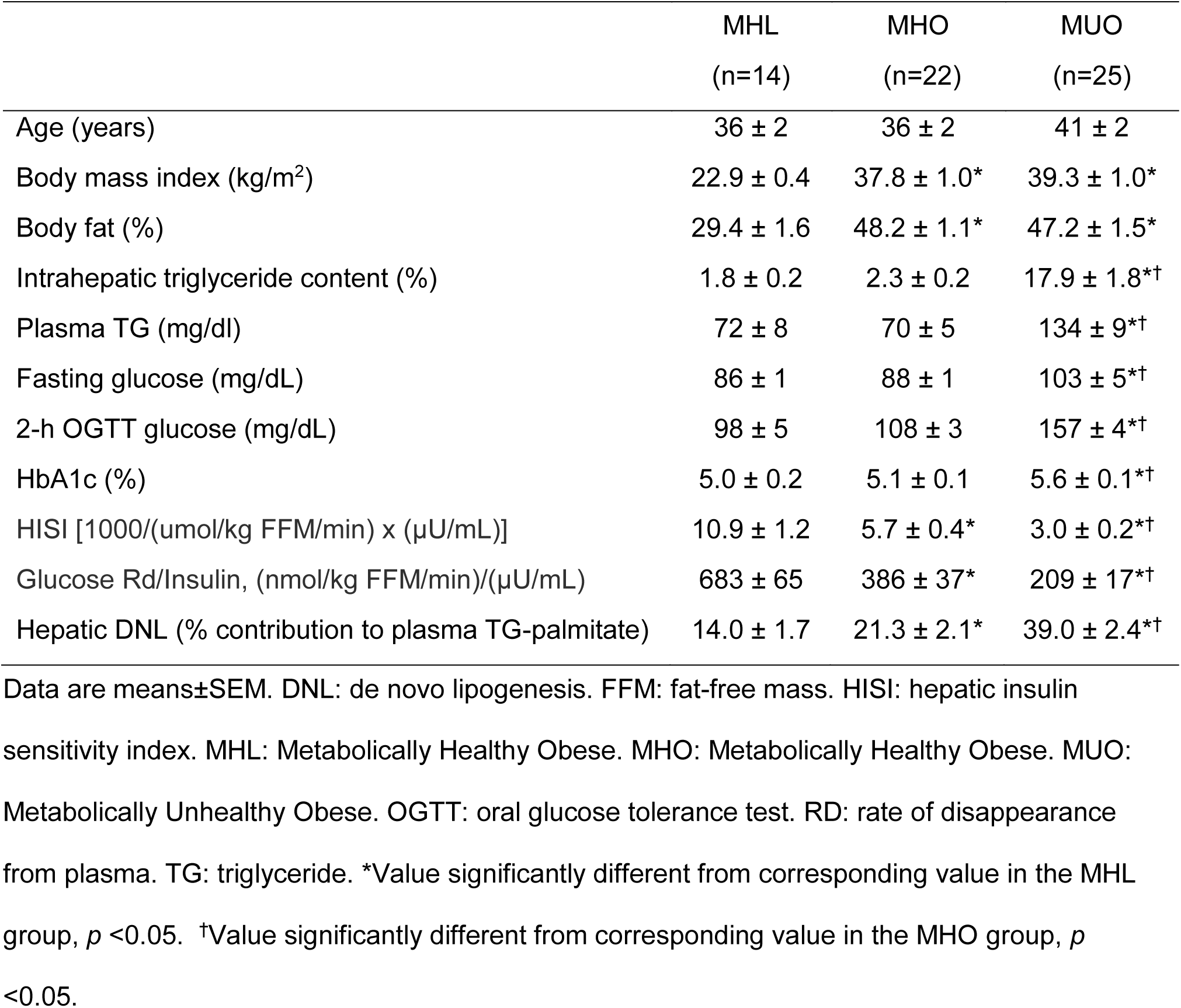
Subject Characteristics.

|  | MHL<br>(n=14) | MHO<br>(n=22) | MUO<br>(n=25) |
| --- | --- | --- | --- |
| Age (years) | 36 ± 2 | 36 ± 2 | 41 ± 2 |
| Body mass index (kg/m<sup>2</sup>) | 22.9 ± 0.4 | 37.8 ± 1.0* | 39.3 ± 1.0* |
| Body fat (%) | 29.4 ± 1.6 | 48.2 ± 1.1* | 47.2 ± 1.5* |
| Intrahepatic triglyceride content (%) | 1.8 ± 0.2 | 2.3 ± 0.2 | 17.9 ± 1.8*† |
| Plasma TG (mg/dl) | 72 ± 8 | 70 ± 5 | 134 ± 9*† |
| Fasting glucose (mg/dL) | 86 ± 1 | 88 ± 1 | 103 ± 5*† |
| 2-h OGTT glucose (mg/dL) | 98 ± 5 | 108 ± 3 | 157 ± 4*† |
| HbA1c (%) | 5.0 ± 0.2 | 5.1 ± 0.1 | 5.6 ± 0.1*† |
| HISI [1000/(umol/kg FFM/min) x (μU/mL)] | 10.9 ± 1.2 | 5.7 ± 0.4* | 3.0 ± 0.2*† |
| Glucose Rd/Insulin, (nmol/kg FFM/min)/(μU/mL) | 683 ± 65 | 386 ± 37* | 209 ± 17*† |
| Hepatic DNL (% contribution to plasma TG-palmitate) | 14.0 ± 1.7 | 21.3 ± 2.1* | 39.0 ± 2.4*† |

Data are means±SEM. DNL: de novo lipogenesis. FFM: fat-free mass. HISI: hepatic insulin sensitivity index. MHL: Metabolically Healthy Obese. MHO: Metabolically Healthy Obese. MUO: Metabolically Unhealthy Obese. OGTT: oral glucose tolerance test. RD: rate of disappearance from plasma. TG: triglyceride. \*Value significantly different from corresponding value in the MHL group, $p < 0.05$ . †Value significantly different from corresponding value in the MHO group, $p < 0.05$ .

**Fig. 1:**
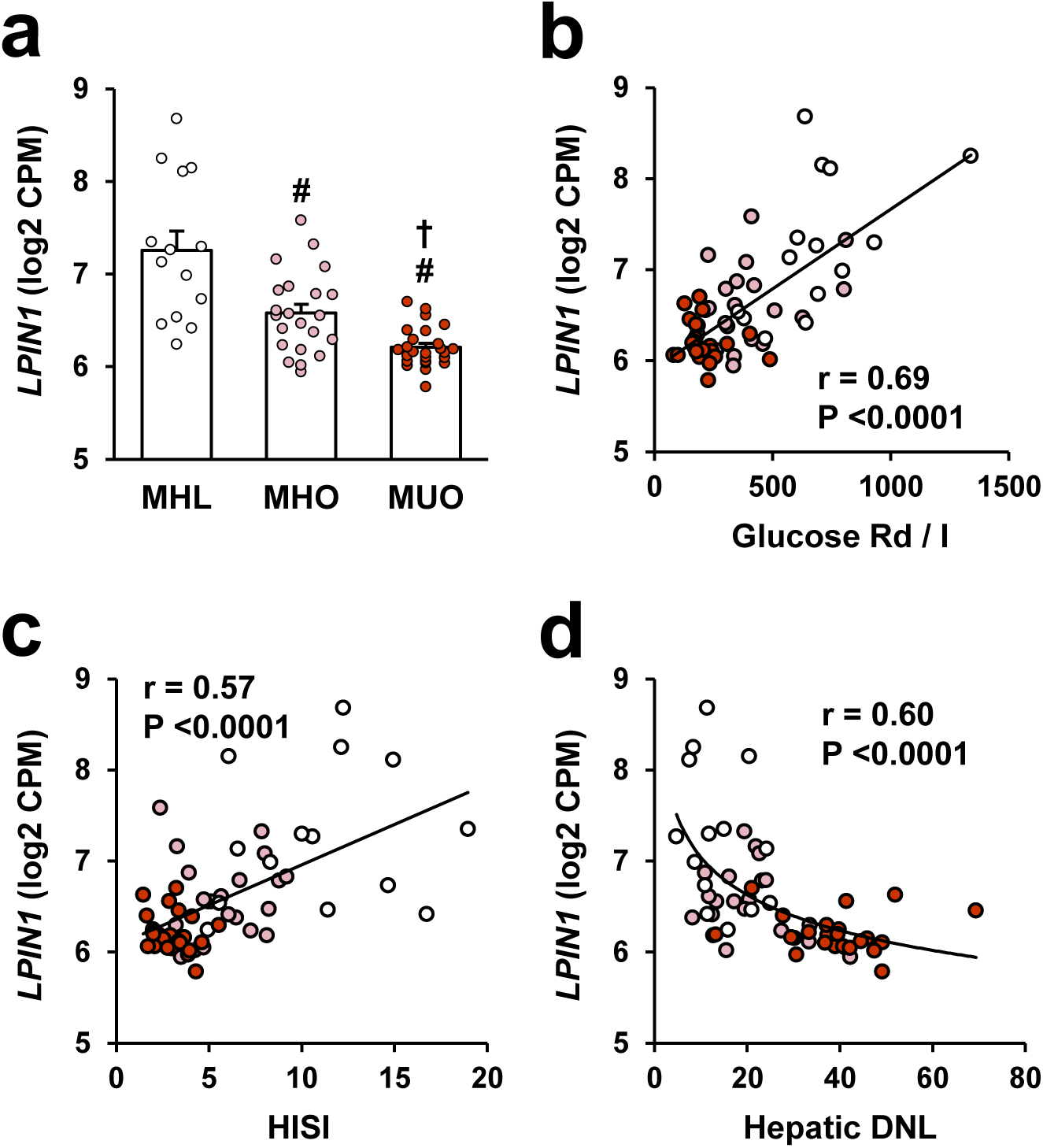
Abdominal adipose tissue *LPIN1* expression is decreased in people with metabolically unhealthy obesity and *LPIN1* expression correlates with metabolic health. **a,** Gene expression of *LPIN1* from subcutaneous abdominal adipose tissue (SAAT) determined by RNA-sequencing in metabolically healthy lean (MHL; n=14), metabolically healthy obese (MHO; n=22) and metabolically unhealthy obesity (MUO; n=25) participants. Data are expressed as means ± SEM. One-way analysis of variance was used to compare *LPIN1* expression among MHL, MHO and MUO groups with Fisher’s least significant difference post-hoc procedure used to identify significant mean differences. ^#^*p* < 0.05 vs. MHL and ^†^*p* < 0.05 vs. MUO. **b-d** Relationship between SAAT *LPIN1* expression and hepatic insulin sensitivity index (HISI), skeletal muscle insulin sensitivity (glucose rate of disappearance relative to plasma insulin concentration during the hyperinsulinemic-euglycemic clamp procedure [glucose Rd/I]) and contribution from hepatic de novo lipogenesis (DNL) to plasma triglyceride-palmitate.

### Adipocyte-specific *Lpin1* knockout mice have reduced fat mass

To test the consequences of *Lpin1* loss in adipose tissue on systemic metabolism, we generated mice with adipocyte-specific *Lpin1* deletion (Adn-*Lpin1-/-*) by using *Lpin1* floxed mice crossed with mice expressing adiponectin promoter-driven Cre recombinase. Prior work to delete *Lpin1* in adipocytes used a distinct line of floxed mice that resulted in expression of a truncated, partially functional protein^21,22^. However, in the present study we used mice with a floxed allele shown to result in complete loss of lipin 1 protein in skeletal muscle and heart^27,28^. Loss of lipin 1 protein and *Lpin1* gene expression was restricted to adipose tissue (Extended Data Fig. 1a,b). There was an increase in lipin 2 mRNA abundance and protein expression in adipose tissue of the Adn-*Lpin1-/-* mice (Extended Data Fig. 1a,b), though previous reports have indicated that the contribution of lipin 2 to adipose tissue PAP activity is minimal^17,32^. At eight weeks of age, male and female Adn-*Lpin1-/-* mice had similar body weights and body composition on chow diet compared to littermate wild-type (WT) control mice (*Lpin1* floxed) (Extended Data Fig. 2). Although female knockout mice were mildly hyperglycemic compared to WT controls, neither sex exhibited differences in an insulin tolerance test (ITT, Extended Data Fig. 2).

Next, we fed male mice a high-fat diet (HFD, 60% kcal from fat) or a matched low-fat diet (LFD, 10% kcal from fat) to determine the effects of caloric excess on the phenotype. After 5 weeks of diet, WT mice fed HFD had a significant increase in body weight, gonadal and inguinal white adipose tissue (gWAT and iWAT, respectively) mass, and percent adiposity compared to WT mice on LFD, while mice lacking adipocyte *Lpin1* gained less weight on the HFD and exhibited reduced fat pad weights compared to WT comparators on either diet (Fig. 2a-e). Adipose tissue from Adn-*Lpin1*-/- mice on both diets exhibited decreased expression of *Pparg1* and adiponectin (*Adipoq*) and increased expression of collagen type 1 alpha (*Col1a1*), transforming growth factor beta 1 (*Tgfb1*), and CD68 (*Cd68;* a macrophage marker) compared to WT control mice (Fig. 2f-o). Gross histology of gWAT and iWAT revealed increases in fibrosis and “crown-like” inflammatory structures in Adn-*Lpin1*-/- mice compared to WT mice and this was exacerbated by HFD feeding (Fig 2p,q).

**Fig. 2:**
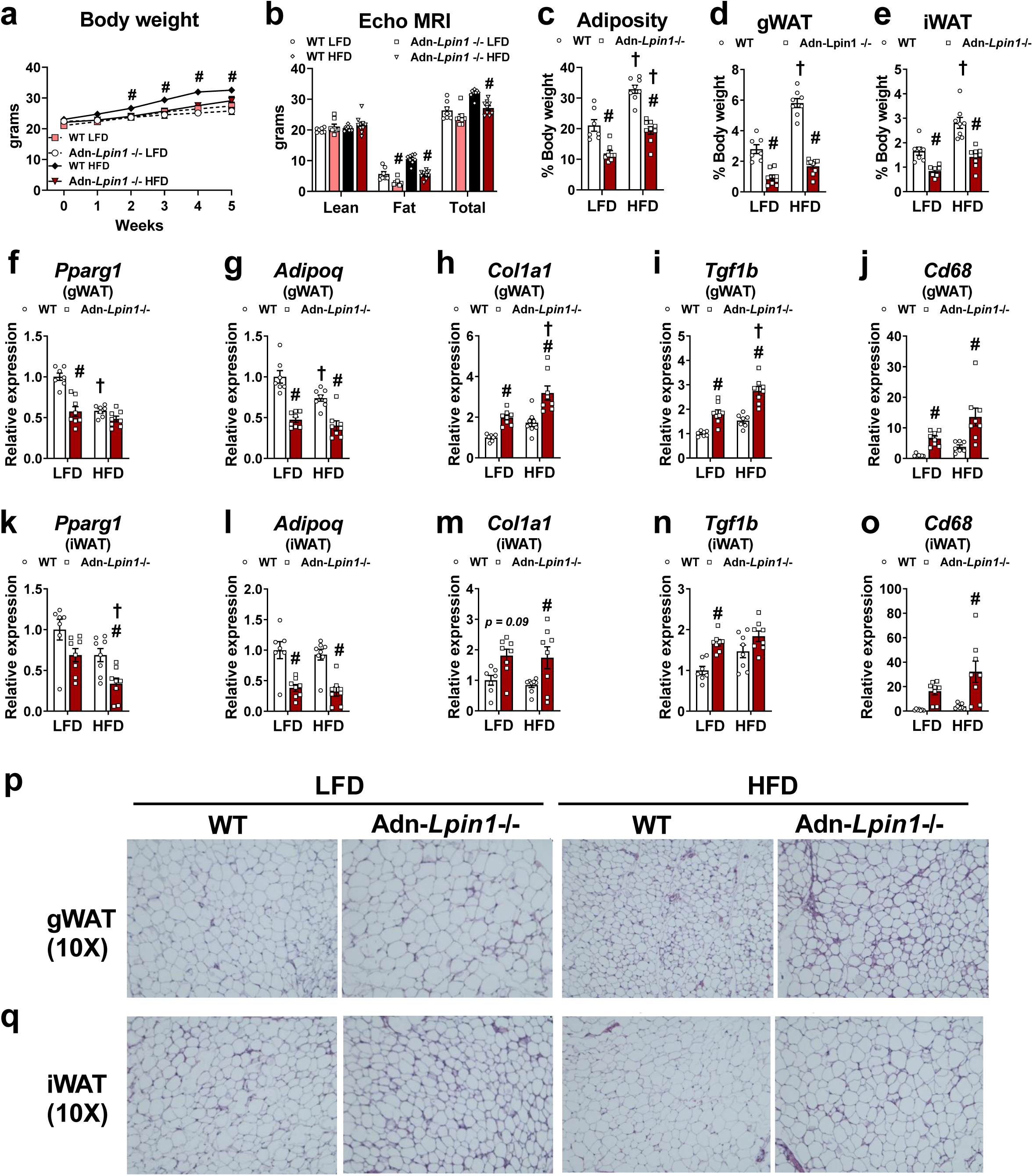
Adipocyte-specific *Lpin1* knockout mice have reduced adiposity and signs of adipose tissue dysfunction. Eight-week-old male adipocyte-specific *Lpin1* knockout mice (Adn-*Lpin1-/-*) and their wild-type littermate controls (WT) were fed either a 10% low-fat diet (LFD) or a 60% high-fat diet (HFD) for 5 weeks. Mice were fasted for 4 h prior to sacrifice and tissue collection. **a,** Body weights of mice during diet feeding. **b,** Body composition was measured in fed mice after 5 weeks of diet via ECHO MRI. **c,** Adiposity was calculated as % fat mass / total body weight. **d,e,** Tissue weight of gonadal white adipose tissue (gWAT) and inguinal white adipose tissue (iWAT) expressed as % body weight. **f-o,** gWAT and iWAT gene expression was determined by qPCR and are expressed as relative abundance; peroxisome proliferator-activated receptor gamma 1 (*Pparg1*), adiponectin (*Adipoq*), collagen type I alpha 1 chain (*Col1a1*), transforming growth factor beta 1 (*Tgfb1*), and cluster of differentiation 68 (*cd68*). **p,q,** Representative 10X magnification images of gWAT and iWAT that were fixed in formalin prior to paraffin embedding, sectioning, and staining with hematoxylin and eosin (H&E). Data are expressed as means ± S.E.M., and significance was determined by Two-way ANOVA with post-hoc Tukey’s multiple comparisons tests. ^#^*p* < 0.05 for WT vs. Adn-*Lpin1-/-* and ^†^*p* < 0.05 for LFD vs. HFD; (*n* = 5-9).

Adipose tissue influences systemic metabolism through the regulated release of non-esterified fatty acids (NEFA), glycerol, and a variety of metabolic peptide hormones known as adipokines. In support of previous findings^21^, mice lacking lipin 1 exhibited reduced plasma NEFA and glycerol concentrations, while TAG concentrations were unaffected by *Lpin1* loss (Extended Data Fig 3a-c). Plasma adiponectin concentrations were significantly reduced in Adn- *Lpin1-/-* mice compared to WT mice on both diets (Extended Data Fig. 3d). Plasma leptin and resistin concentrations tended to be lower in knockout mice but were not significantly reduced (Extended Data Fig. 3f). Together these data suggest that loss of adipocyte lipin 1 is sufficient to cause adipose tissue dysfunction in mice.

### Loss of adipocyte *Lpin1* leads to increased respiratory exchange ratio and food consumption in LFD fed mice, but this is corrected by HFD feeding

Given that Adn-*Lpin1-/-* mice are leaner than their WT littermates, we utilized a metabolic cage system to define metabolic rates, substrate preferences, activity, and food consumption. On the LFD, Adn-*Lpin1-/-* mice exhibited a non-significant trend towards increased energy expenditure, a significantly higher respiratory exchange ratio (RER) and cumulative food intake, with no change in ambulatory activity (Fig 3a-d). These effects were present in both the light and dark phases. The higher RER indicated increased rates of carbohydrate metabolism versus fat oxidation, which is consistent with reduced circulating NEFA in Adn-*Lpin1-/-* mice (Extended Data Fig 3 a). Importantly, analysis of the data using CalR,^33^ demonstrated a significant interaction between O_2_ consumption and body weights, suggesting that some of these effects are driven by differences in body composition. However, no difference in energy balance between genotypes was detected (data not shown). We also assessed these parameters in mice on the HFD and found that high dietary fat content abrogated the trend towards increased energy expenditure and genotype effects on RER and food consumption (Fig. 3e-h). This suggests that dietary fat abolishes the *Lpin1* genotype effect on RER by increasing fat availability to normalize energy substrate utilization.

**Fig 3:**
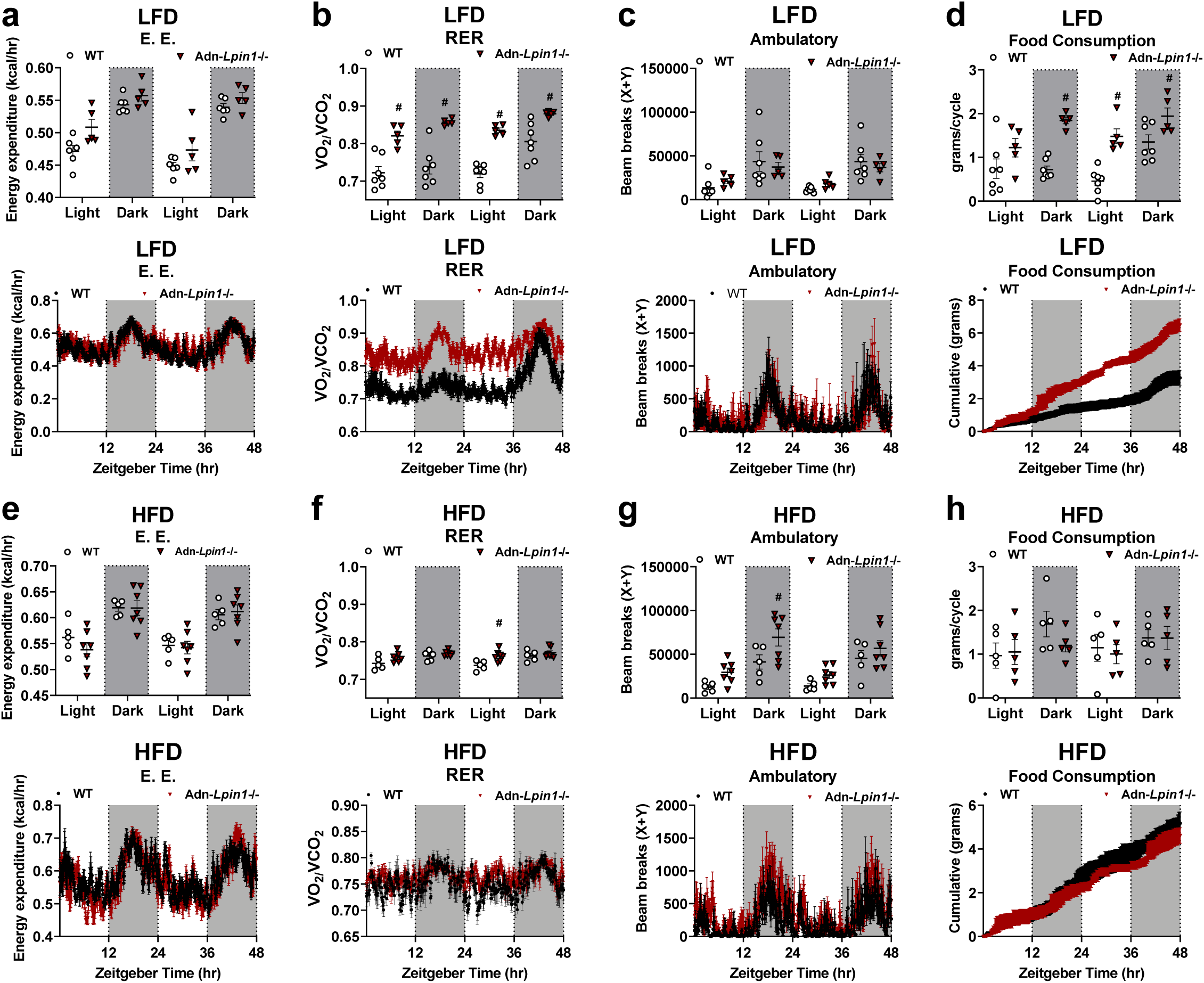
Adn-*Lpin1*-/- mice utilize consume and utilize more carbohydrates on a LFD compared to control mice. Eight-week old male mice were fed a LFD or HFD or 5 weeks prior to body composition analysis via EchoMRI and single-housing in a 16-metabolic cage Comprehensive Laboratory Animal Monitoring System (CLAMS, Columbus Instruments). **a-d,** Energy expenditure (E.E), respiratory exchange ratio (RER), ambulatory movement, and food consumption were averaged for each 12 hour cycle (top) or shown as measured over 48 hours (bottom) for mice fed a 10% LFD (*n* = 5-7). **e-h,** The same measurements were conducted for mice fed a 60% HFD (*n* = 5-7). Data are expressed as means ± S.E.M., and significance was determined by Two-way ANOVA with post-hoc Tukey’s multiple comparisons tests. ^#^*p* < 0.05 for WT vs. Adn-*Lpin1-/-*.

### Adn-*Lpin1-/-* mice are metabolically unhealthy

Insulin and glucose tolerance tests were performed with WT and Adn-*Lpin1*-/- mice on either LFD or HFD. As observed with younger mice on a chow diet, 12-13 week old Adn-*Lpin1-/-* mice fed a LFD exhibited similar area under the curve (AUC) values during ITT (week 4 of diet) and GTT (week 5 of diet) analyses compared to WT mice (Extended Data Fig.4). In contrast, Adn-*Lpin1-/-* mice fed a HFD had higher fasting glucose concentrations compared to all other groups and were significantly insulin and glucose intolerant compared to all other mice after HFD feeding (Extended Data Fig. 4). In contrast, this short-term HFD treatment did not induce glucose or insulin intolerance in HFD-fed WT mice compared to LFD-fed WT mice.

WT and Adn-*Lpin1-/-* mice fed a HFD underwent hyperinsulinemic-euglycemic clamp studies as the gold standard for insulin sensitivity and to determine which tissues were insulin-resistant (Fig. 4). Adn-*Lpin1* -/- mice were hyperglycemic at the start of the clamp procedure and required significantly lower rates of exogenous glucose infusion to reach steady-state arterial blood glucose concentrations compared to WT mice (Fig. 4a,b). Plasma insulin concentrations were also elevated in the fasting state and during the clamp in Adn-*Lpin1-/-* mice (Fig. 4c). Fasting endogenous glucose production was elevated in Adn-*Lpin1-/-* mice, compared to WT mice, and insulin was less effective at suppressing glucose production during the clamp (Fig. 4d). Considering the rate of glucose appearance as a function of plasma insulin concentrations, displayed as the slope from fasted to clamp states, revealed a significant difference between genotypes indicating that insulin was less effective at suppressing endogenous glucose production and is further evidence of hepatic insulin resistance in Adn-*Lpin1-/-* mice (Fig. 4e). Similarly, compared to WT mice, high insulin concentrations were less effective at increasing the glucose disposal rate in Adn-*Lpin1-/-* mice (Fig. 4f,g). Quantification of 2-deoxyglucose uptake into various tissues revealed that Adn-*Lpin1-/-* mice exhibited a substantial decrease in glucose disposal in subscapular brown adipose tissue (BAT) and vastus muscle, with a trend towards reduced uptake in soleus and gastrocnemius muscle (Fig. 4h,i). Collectively, these data suggest that loss of lipin 1 in adipocytes leads to systemic insulin resistance that impacts liver, skeletal muscle, and BAT.

**Fig. 4:**
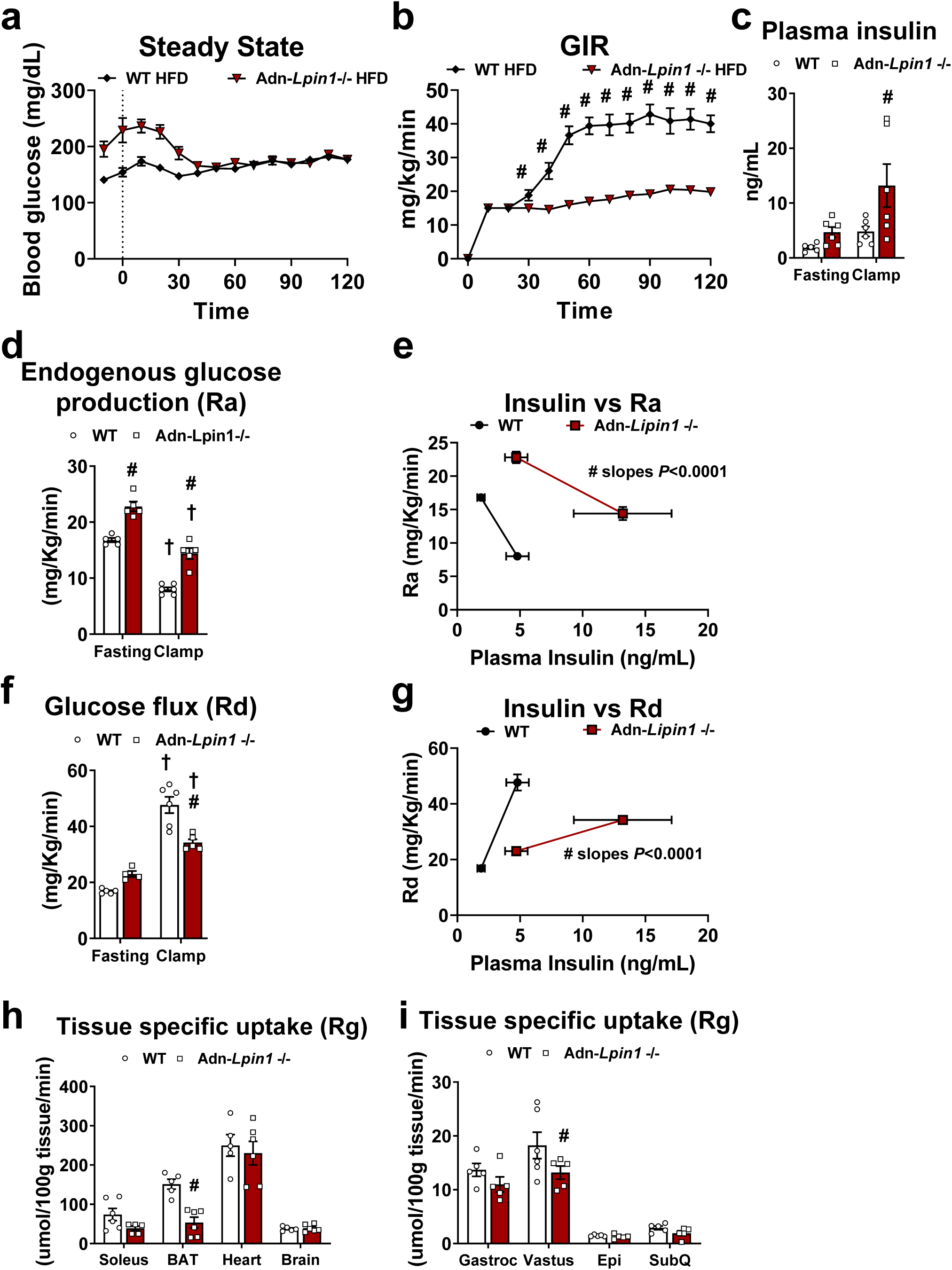
Adn-*Lpin1-/-* mice exhibit systemic insulin resistance on HFD. Eight-week-old male Adn-*Lpin1-/-* and WT control mice were fed a 60% HFD for 5 weeks. Five days prior to clamp mice were catheterized and allowed to recover. Mice were fasted 5 hours prior to the clamp procedures as described in detail in the methods section. **a,** Blood glucose was monitored prior to and during the clamp and shows both groups reached and sustained similar steady state glucose concentrations during the clamp procedure. **b,** Exogenous glucose infusion rates (GIR) were measured during the clamp procedure. **c,** Plasma insulin was measured by radioimmunoassay at -10 min for fasting and 90 and 120 minutes were averaged for clamp values. **d,** Endogenous glucose rate of appearance (Ra) was determined from steady-state equations. **e,** Ra was plotted against plasma insulin concentrations prior to and during the clamp. **f,** Whole-body glucose disposal (Rd) was determined from steady-state equations. **g,** Rd was plotted against plasma insulin concentrations prior to and during the clamp. **h,I,** Tissue-specific glucose uptake (Rg). Data are expressed as means +/- standard error of the means (S.E.M), and significance was determined by Student’s T test or two-way ANOVA with post-hoc Tukey’s multiple comparisons tests where appropriate. ^#^*p* < 0.05 for WT vs Adn-*Lpin1-/-* and ^†^*p* < 0.05 for LFD vs HFD; (*n* = 5-6).

### Loss of adipocyte *Lpin1* leads to ectopic lipid accumulation and increased hepatic *de novo lipogenesis*

On both the LFD and HFD diets, Adn-*Lpin1-/-* mouse livers were enlarged compared to WT control mice and appeared to accumulate lipids as indicated by the vacuolar appearance of sections after H&E staining (Fig. 5a,b). To quantify the abundance of various lipid species we performed targeted lipidomic analyses. We found that compared to WT mice, intrahepatic DAG and TAG was much higher (10- and 5-fold, respectively) in Adn-*Lpin1-/-* mice fed LFD (Fig 5c). While HFD increased hepatic glycerolipid content in WT mice, the effect of HFD was more dramatic in Adn-*Lpin1-/-* mice compared to WT mice (Fig. 5c). Several species of TAG with long polyunsaturated acyl chains were significantly increased, most notably 20:4 (Fig. 5c). Similar increases in DAG, but not TAG, were observed in lipidomic analyses of plasma and gastrocnemius of these mice (Extended Data Fig. 5). Given that Adn-*Lpin1-/-* mice exhibited lower plasma NEFAs and were insulin resistant, which is known to drive increased hepatic DNL^34^, we sought to determine rates of DNL in Adn-*Lpin1-/-* mice on HFD. Indeed, isotopic tracing studies conducted using deuterated water confirmed an increase in the fractional synthesis rate of liver palmitate in Adn-*Lpin1-/-* mice compared to WT mice on HFD (Fig 5d). These data indicate that loss of *Lpin1* in adipocytes leads to hepatic steatosis in conjunction with increased rates of DNL.

**Fig. 5:**
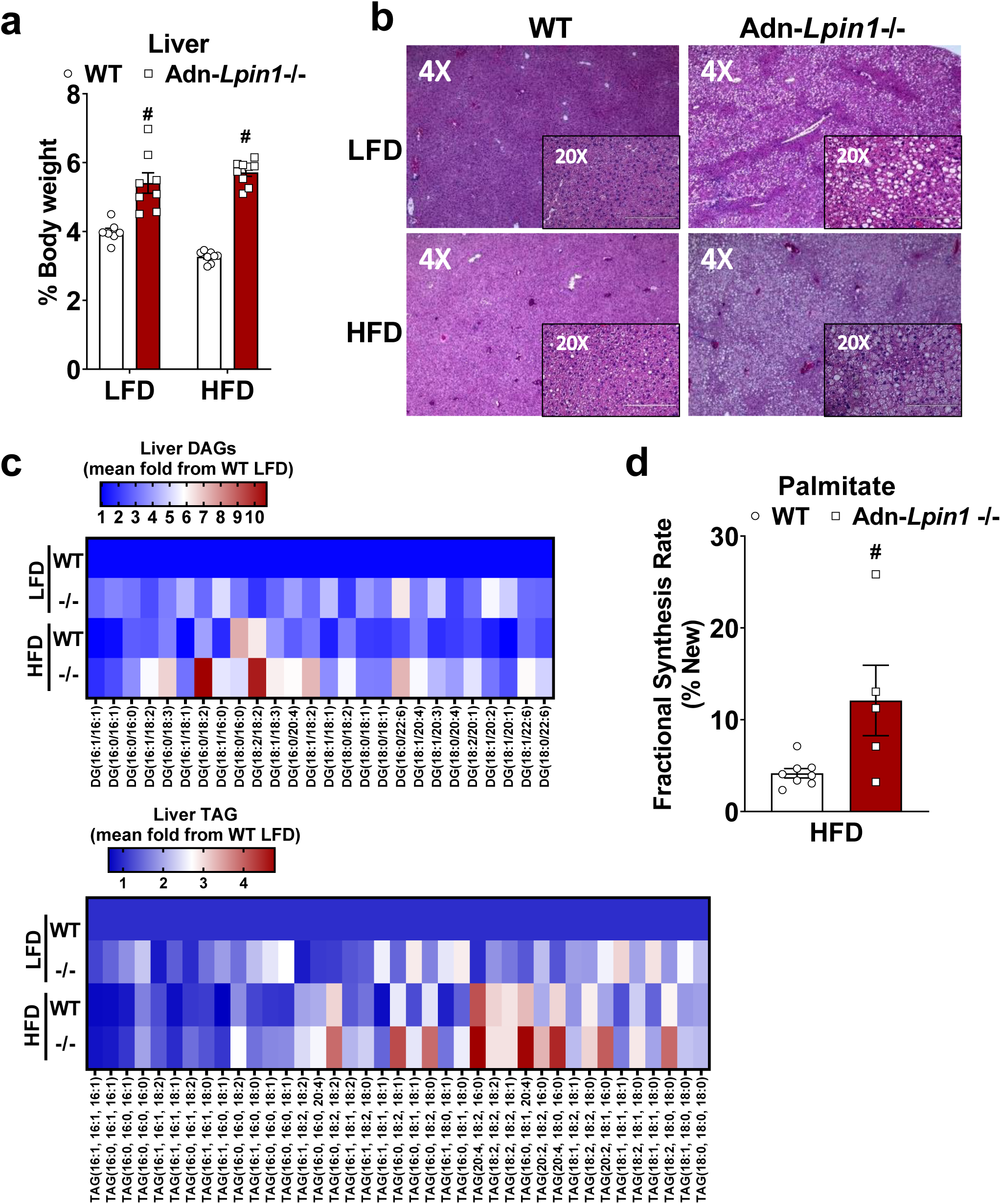
Loss of adipocyte *Lpin1* leads to severe hepatic lipid accumulation. Eight-week-old male Adn-*Lpin1-/-* and WT control mice were fed either a 10% LFD or a 60% HFD for 5 weeks Mice were fasted for 4 hours prior to sacrifice and tissue collection. **a,** Liver weight expressed as % of body weight. **b,** Representative 4X and 20X magnification images of liver tissue that were fixed in formalin prior to paraffin embedding, sectioning, and staining with H&E. **c,** Liver diacylglycerols (DAG) and triglycerides (TAG) were extracted and the relative abundance of each species was determined by LC-MS/MS against an internal standard. Data are expressed as mean fold change from the Control LFD-fed mice; (*n* = 7-9). **d,** Total palmitate was analyzed by LC-MS for isotope mass abundance and MIDA was used to calculate palmitate fractional synthesis rate (% newly synthesize). Data are expressed as means +/- S.E.M, and significance was determined by a Student’s T-test. ^#^*p* < 0.05 for WT vs. Adn-*Lpin1-/-;* (*n* = 5-8).

### Bulk RNA sequencing analysis reveals significant changes in the expression of genes in pathways related to fatty liver to NASH transition

To identify the molecular pathways that are associated with insulin resistance and steatosis in Adn-*Lpin1-/-* mice, we performed bulk RNA sequencing in liver (Fig. 6 and Extended Data Fig. 6). PCA analyses demonstrated significant separation among the four groups with apparent gene changes represented by a heat map. (Extended Data Fig. 6a,b). There was a significant increase of genes associated with hepatic steatosis (*Cidea*, *Cidec*, and *Plin4*) and fibrosis (*Col1a1*) in Adn-*Lpin1-/-* livers on both diets compared to their diet-matched control mice (Fig. 6a,b). Next, we performed weighted gene co-expression analysis (WGNCA) to identify highly correlated gene modules within the sequencing data (Fig. 6c) ^35^. Several modules significantly correlated with the genotype of the mice and/or the diets (Figure 6c, select modules shown). The turquoise module contained the largest number of genes at 2762 and most of the significantly upregulated genes in the volcano plots clustered to this module (Fig. 6a,b and Extended Data Table 1). Additionally, the turquoise module was positively correlated to the HFD fed Adn-*Lpin1-/-* mice and most negatively with the control LFD fed mice (Fig. 6c).

**Fig. 6:**
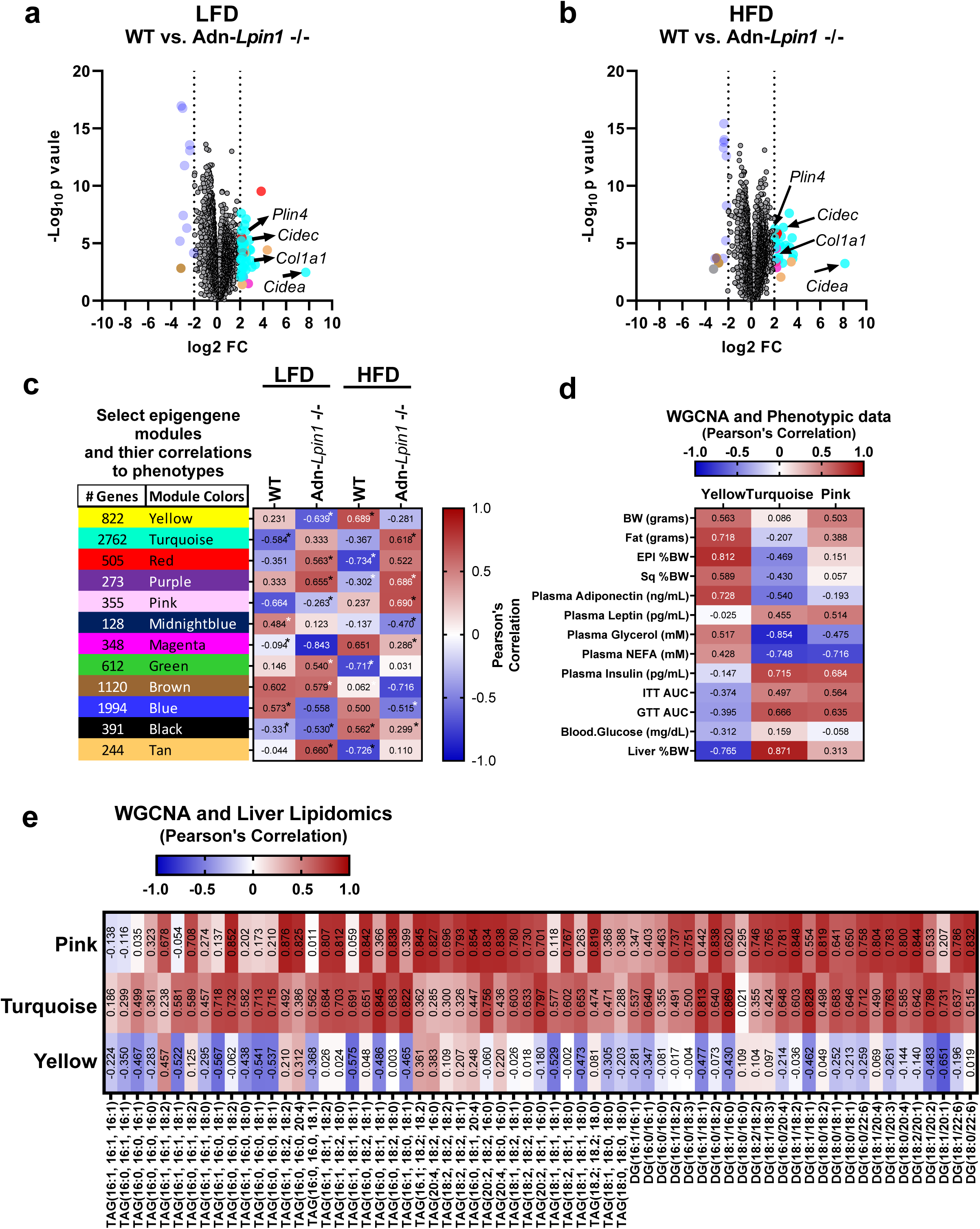
Bulk RNA sequencing analysis reveals significant changes in genetic pathways related to early-stage fatty liver to NASH transition. Eight-week-old male Adn-*Lpin1-/-* and WT control mice were fed either a 10% LFD or a 60% HFD for 5 weeks. Mice were fasted for 4 hours prior to sacrifice, liver collection, RNA isolation, and Bulk RNA sequencing. **a,b,** Volcano plots of merged differential expression data were graphed as log_2_ fold change versed –log_10_ unadjusted *p-*value. The color of the data points corresponds to the module eigengenes each gene clusters to based on hierarchical clustering. **c,** Select gene modules and the number of genes in each module from weighted gene co-expression analysis (WGCNA). **d,** Signification modules and their correlation to DAG and TAG species and e, phenotypic traits from the mice used to generate the WGCNA data. Pearson’s Correlation Coefficient was used to determine association with the WGCNA modules and trait data; \**p* < 0.05, (*n* = 6).

We next combined WGCNA module sets with phenotypic traits from the mice used to generate the RNA sequencing. Using Pearson’s Correlation Coefficient, we found several modules sets that were significantly correlated to the phenotypic data such as tissue size, plasma adiponectin, glycerol, and NEFAs (Fig. 6d and Extended Data Fig. 7a). In addition, we compared our lipidomic data to identify gene modules that correlate with the increased DAG and TAG species seen in Adn-*Lpin1-/-* livers and discovered that many of these lipid species also had significant correlations to gene modules (Fig. 6e and Extended Data Fig. 7b). Specifically, turquoise and pink modules positively correlated with plasma insulin levels, liver size, and most DAG and TAG species. Alternatively, these modules were negatively correlated to plasma glycerol and NEFAs (Fig. 6d,e and Extended Data Fig. 7). Pathway analysis of the turquoise module genes revealed several significant GO Molecular pathways involved in extracellular matrix components, lipid binding, and collagen binding (Extended Data Fig. 8). In fact, one of the most significantly upregulated genes in Adn-*Lpin1-/-* livers is *Col1a1*, which clusters to the turquoise model and is a marker of activated stellate cells, the primary mediators of hepatic fibrosis (Fig 6a,b and Extended Data Table 1). Significant KEGG pathways within the turquoise module included PI3K-Akt signaling, ECM-Receptor interactions, and AGE-RAGE signaling pathway in diabetic complications (Extended Data Fig. 8a). Pathway analysis of the pink module set included significant GO molecular function pathways involved in oxidoreductase activity FAD/NAD binding and significant GO biological pathways involved in metabolic and catabolic pathways specifically related to lipid metabolism (Extended Data Fig. 8b). Lastly, the yellow module, which most negatively correlated to Adn-*Lpin1-/-* mice had positive correlations for fat mass, adiponectin, plasma glycerol and NEFA and negative correlations for ITT AUC, GTT AUC, blood glucose, and liver size. Yellow pathway analysis revealed significant changes in FoxO, mTOR, AMPK, and insulin signaling pathways, as well as regulation of kinase signaling and activity (Extended Data Fig. 8c). Because insulin resistance and liver disease progression are highly related^36^ we confirmed the increased expression of several markers of non-alcoholic steatohepatitis (NASH) development in Adn-*Lpin1-/-* mice by qRT-PCR. We demonstrated increased expression of genes related to fibrosis (*Col1a1*), matrix remodeling (*Timp1* and *Timp3*), inflammation (*Spp1*, *Cd68*, *Il1b*), and *Tgfb1* compared to WT mice on the same diet (Extended Data Fig. 9a). Together these data suggest that loss of adipocyte lipin1 drives insulin resistance and increases early markers of hepatic fibrosis and NASH.

### Loss of adipocyte-*Lpin1* predisposes mice to liver injury and hepatic stellate cell activation

Although Adn-*Lpin1-/-* mice exhibited transcriptional profiles consistent with the development of NASH, there were no significant changes in plasma ALT or AST levels, surrogate markers of liver damage, and histological scoring of H&E liver sections did not reveal any significant changes (Extended Data Fig. 9b,c). Because mice are inherently resistant to developing NASH on standard high-fat diets^37^, we used a diet high in fructose (17 kcal %), fat (mostly palm oil 40 kcal %), and cholesterol (2%) (HFHF-C) or a matched sucrose, high sugar (dextrose), low-fat (10 kcal % fat) control diet (HSLF) (Fig. 7). After 16 weeks of diet, the HFHF- C diet did not cause weight gain or changes in blood glucose or plasma insulin concentrations in mice fed the HFHF-C diet (Fig 7a and Extended Data Fig. 10a). However, Adn-*Lpin1*-/- mice were leaner on both diets compared to WT mice (Fig. 7a-d). Analogous to the short-term HFD studies (Extended Fig. 3), Adn-*Lpin1*-/- mice had similar changes in plasma concentrations of NEFA, glycerol, and TAG while cholesterol was increased compared to WT mice (Extended Data Fig. 10b). The livers of Adn-*Lpin1-/-* mice were larger compared to controls reaching about 13% of body weight on the HFHF-C diet (Fig. 7e). Adn-*Lpin1*-/- mice also had enlarged spleens and significant elevations in plasma ALT, but not AST, on the HFHF-C diet indicating increased liver injury (Fig. 7f-h). There was no significant difference in hepatic TAG content between genotypes and only trends towards an increase in NAFLD scoring (Fig. 7i-l). However, gene expression markers of hepatic stellate cell activation and inflammation were increased in Adn-*Lpin1*-/- on HFHF-C diet compared to WT control mice (Fig. 7m-o). These data indicate that the loss of adipocyte *Lpin1* exacerbates the liver injury and activation of markers of NASH in mice.

**Fig. 7:**
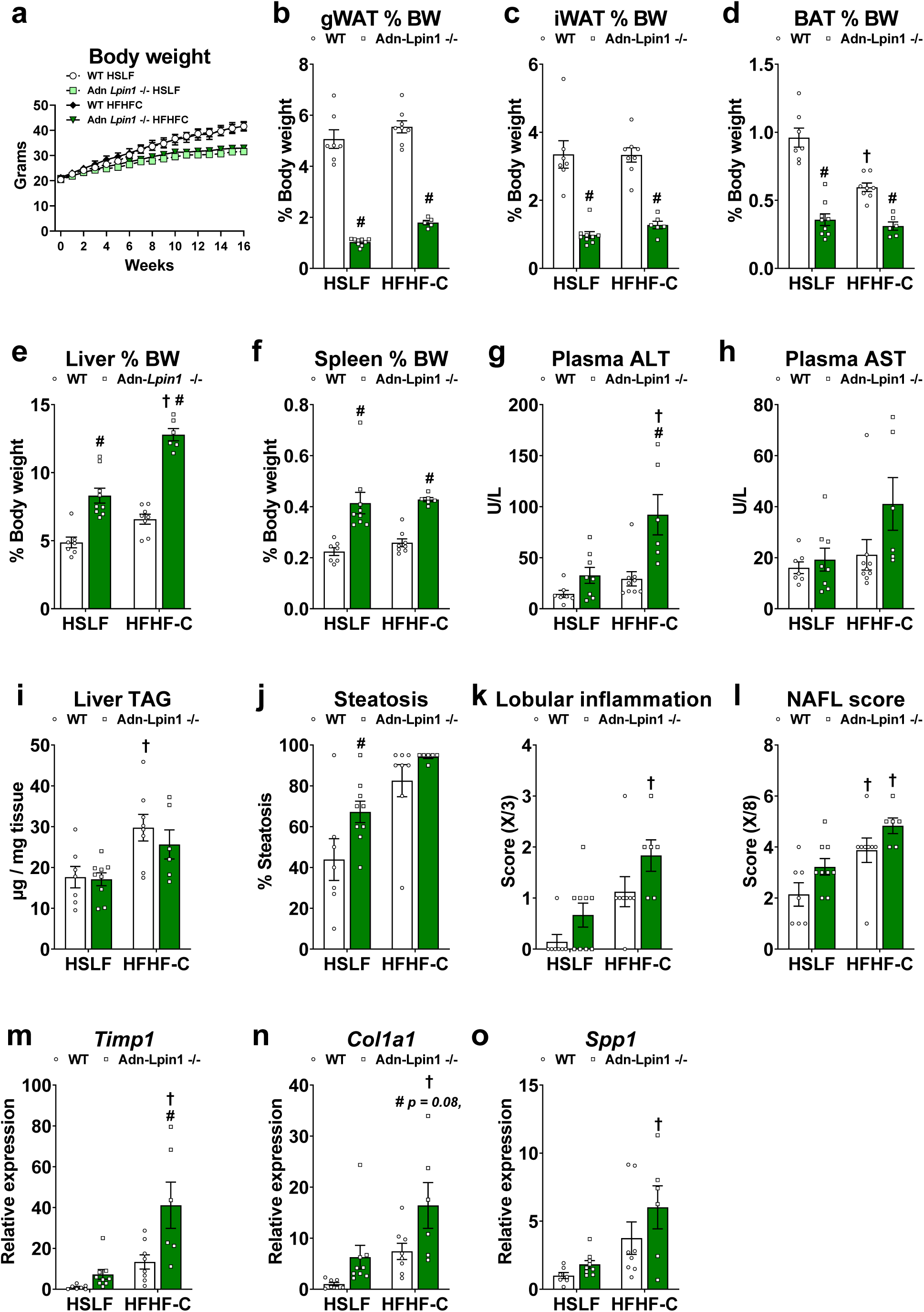
Loss of adipocyte-*Lpin1* predisposes mice towards NASH. Eight-week-old male mice were fed a diet high in fructose (17 kcal %), fat (mostly palm oil 40 kcal %), and cholesterol (2%) (HFHF-C) or a matched sucrose, high sugar (dextrose), low-fat (10 kcal % fat) control diet (HSLF) for 16 weeks. Mice were fasted for 4 hours prior to sacrifice and tissue collection. **a,** Weekly body weights. **b-e,** Individual tissue weights expressed as % total body weight. **g,h,** Plasma alanine transferase (ALT) and aspartate aminotransferase (AST) were measured using liquid kinetic assays. **i,** Liver lipids were extracted and quantified using a colorimetric enzymatic assay. **j-l,** H&E stained tissue sections were scored by an independent clinical pathologist. **e-g,** Gene expression was determined by qPCR and are expressed as relative abundance; tissue inhibitor of metalloproteinase 1 (*Timp1*), collagen type I alpha 1 chain (*Col1a1*), secreted phosphoprotein 1 (*Spp1*). Data are expressed as means +/- S.E.M, and significance was determined by Two-way ANOVA and post-hoc Tukey’s or Sidak’s multiple comparisons tests. ^#^*p* < 0.05 for WT vs. Adn-*Lpin1-/-* and ^†^*p* < 0.05 for HSLF vs HFHF-C diet; (*n* = 7-9).

## Discussion

Dysfunctional adipose tissue may play a critical role in the development of metabolic abnormalities and diseases, including insulin resistance and NAFLD. The mechanisms by which this occurs remain incompletely understood. Herein, we have shown that adipocyte *LPIN1* expression is reduced in people with obesity compared to people who are MHL and its expression correlates with measures of skeletal muscle and hepatic insulin sensitivity and inversely correlates with hepatic de novo lipogenesis, a sensitive marker of hepatic insulin resistance and a characteristic of NAFLD^34^. In mice, the loss of *Lpin1* in adipocytes leads to a lean phenotype, but promotes adipose tissue dysfunction and negatively affects systemic metabolism. Although Adn-*Lpin1-/-* mice were lean, they showed features of insulin resistance and steatosis observed in our participants. A recent study suggests that increased BMI and visceral adiposity are associated with increased adipose tissue inflammation and NAFLD, even in lean subjects, and tracks with the severity of liver disease^38^.These data are consistent with the idea that systemic influences of adipose tissue on metabolism are based on quality, rather than quantity of adipose tissue. Dysfunctional adipose tissue can be defined by a variety of qualities including diminished capacity for lipid storage, insulin resistance, increased inflammation and fibrosis, and altered secretion of adipokines and other factors^9,39–41^. Deletion of *Lpin1* in adipocytes impacted each of these parameters as we have observed reduced rates of TAG synthesis^21^, signs of adipose tissue inflammation and fibrosis, and reduced secretion of the beneficial adipokine, adiponectin.

One of the most striking phenotypes we observed was the increased hepatic insulin resistance and lipid accumulation (Fig. 4,5) and the demonstration that loss of adipocyte *Lpin1* is sufficient to drive liver injury in mice fed a NASH-inducing diet (Fig. 7). Bulk RNA sequencing of liver tissue and WGCNA analysis revealed that Adn-*Lpin1*-/- mice exhibit dysregulation of several molecular signaling pathways that are known to impact the development of NASH (Yellow Module, Fig. 6 and Extended Data Fig. 8). Moreover, the accumulated intrahepatic DAG/TAG species correlate to increased expression of genes and genetic pathways associated with fatty liver to NASH transition including *Col1a1* and cell adhesion/ECM remodeling pathways (Fig. 6,7 and Extended Data Table 1). The results from our WGCNA analysis reveals strong correlations between DAG/TAG species and these disease pathways (Fig. 6 and Extended Data Fig. 7). Species with long-chain unsaturated fatty acyl chains had the highest correlation with pathways associated redox state, fatty acid metabolism, and NASH (Turquoise and Pink module, Fig 6. and Extended Data Fig. 8). In contrast, on the NASH-inducing diet, the enhanced liver injury exhibited by Adn-*Lpin1-/-* mice was not accompanied by increased intrahepatic TAG content or hyperinsulinemia. This suggests that other factors, including lower circulating adiponectin levels, could also be involved.

Ectopic lipid accumulation in insulin target tissues is tightly linked to the development of hepatic and systemic insulin resistance. In particular, tissue DAG accumulation has been mechanistically linked to the development of liver and skeletal muscle insulin resistance^42^. Prior work had suggested that mice lacking lipin 1 in adipose tissue were insulin-resistant^17,22^, but the specific tissues that are impacted had not been investigated. Hyperinsulinemic clamp studies of HFD-fed Adn-*Lpin1*-/- mice revealed increased rates of glucose appearance from liver and showed that insulin failed to suppress hepatic glucose production. Clamp studies also demonstrated that Adn-*Lpin1*-/- mice exhibited reduced glucose uptake into skeletal muscle and BAT, suggesting that loss of lipin 1 in adipose tissue leads to multi-tissue insulin resistance. Future studies are required to address the specific cause of systemic insulin resistance in mice lacking adipocyte lipin 1.

The sources of the lipids accumulating in liver of Adn-*Lpin1*-/- mice remain to be determined. In this work and prior studies^21^, we show that plasma NEFA and glycerol concentrations are actually lower in adipocyte-specific *Lpin1* knockout mice (Extended Data Fig. 3a,b) suggesting the hepatic lipids are not derived from increased lipolysis. Indeed, we have previously shown that loss of lipin 1 in adipocytes leads to reduced rates of lipolysis due to suppression of protein kinase A signaling^21^. In mice fed the high fat diets, hepatic lipids could be derived from the uptake of dietary lipids, but increased DAG and TAG was also observed in Adn-*Lpin1-/-* mice on diets with low fat content. In people with obesity and NAFLD, intrahepatic *de novo* lipogenesis (DNL) is increased several-fold compared to lean subjects and contributes to intrahepatic and plasma TAG^34,43^. We also observed increased fractional synthesis rates of palmitate in Adn-*Lpin1*-/- mice, consistent with our finding in human subjects of an inverse correlation between adipose tissue Adn-*Lpin1-/-* gene expression and hepatic DNL. It is likely that increased rates of DNL observed in these mice, which may be driven by their insulin resistance, contributes to the observed hepatic steatosis even on chow diet.

In summary, the present study bolsters the argument that adipose tissue *function* rather than quantity is critical for preserving metabolic health. This work unveiled a strong correlation between adipose tissue *LPIN1* expression and hepatic and skeletal muscle insulin sensitivity as well as an inverse correlation with hepatic DNL in people. These findings provide evidence that loss of adipocyte lipin 1 is sufficient to induce systemic metabolism perturbations in mice. The mechanisms of interorgan communication involved in these phenotypes are likely multifactorial. Future work is needed to define which factors plays a causal role in the inter-organ communication between adipose tissue and other important insulin target tissues.

## Methods

### Human Studies

#### Study Subjects

Sixty-one men and women participated in this study that was approved by the Human Research Protection Office of Washington University School of Medicine in St. Louis, MO (ClinicalTrials.gov, NCT02706262). Participants were recruited by using the Volunteers for Health database at Washington University School of Medicine and by advertising in the community. Written, informed consent was obtained from all participants before being enrolled in this study. All participants completed a comprehensive screening evaluation, including a medical history and physical examination, standard blood tests, hemoglobin A1c (HbA1c), an oral glucose tolerance test (OGTT), and assessment of intrahepatic triglyceride (IHTG) content by using MRI to determine eligibility. Subjects were characterized by body weight status and metabolic health into three groups: 1) metabolically-healthy lean (MHL; n=14, 8 women) defined as body mass index (BMI) 18.5-24.9 kg/m^2^, IHTG content <5%, plasma TG concentration <150 mg/dL, normal fasting plasma glucose (<100 mg/dL), normal oral glucose tolerance (plasma glucose, <140 mg/dL 2 hours after ingesting 75 g of glucose), and HbA1c (≤5.6%); 2) metabolically-healthy obese (MHO; n=22, 21 women) defined as BMI 30-49.9 kg/m^2^ with normal IHTG content, plasma TG and glucose tolerance; and 3) metabolically unhealthy obese (MUO; n=9), defined as BMI 30-49.9 kg/m^2^ with IHTG content ≥5.6%, and abnormal glucose metabolism determined by HbA1c 5.7%-6.4%, fasting plasma glucose 100-125 mg/dL, or 2-hour OGTT plasma glucose concentration 140-199 mg/dL. Potential participants who had a history of diabetes or liver disease other than NAFLD, consumed excessive amounts of alcohol (>21 units per week for men and >14 units per week for women), were taking medications that could affect the study outcome measures that precluded MRI were excluded from the study.

### Body composition analyses

Total body fat and fat-free mass (FFM) were determined by using dual-energy X-ray absorptiometry. Intra-abdominal AT (IAAT) and SAAT volumes and IHTG content were assessed by using MRI.

### Insulin sensitivity

Subjects were admitted to the Clinical and Translational Research Unit (CTRU) at Washington University School of Medicine in St. Louis, Missouri at 1700 h for ∼48 hours and consumed standard meals (50% carbohydrate, 35% fat, 15% protein) containing one-third of their estimated energy requirements^44^ at 1800 h on day 1 and at 0700 h, 1300 h, and 1900 h on day 2. After the evening meal was consumed on day 2, participants fasted until the end of the hyperinsulinemic-euglycemic clamp procedure conducted on day 3. At 0700 h on day 3, a primed (8.0 µmol/kg) continuous (0.08 µmol/kg/min) infusion of [U-^13^C] glucose (Cambridge Isotope Laboratories Inc.) was started through the intravenous catheter inserted into an antecubital vein. An additional catheter was inserted into a radial artery to obtain arterial blood samples. After the infusion of glucose tracer for 210 min (basal period), insulin was infused for 210 min at a rate of 50 mU/m^2^ body surface area (BSA)/min (initiated with a two-step priming dose of 200 mU/m^2^ BSA/min for 5 min followed by 100 mU/m^2^ BSA/min for 5 min). The infusion of [U-^13^C]glucose was stopped during insulin infusion because of the expected decrease in hepatic glucose production^45^. Euglycemia (∼100 mg/dl) was maintained by variable infusion of 20% dextrose enriched to ∼1% with [U-^13^C] glucose. Blood samples were obtained before beginning the tracer infusion and every 6-7 minutes during the final 20 minutes (total of 4 blood samples) of the basal and insulin infusion periods. Abdominal subcutaneous adipose tissue was obtained during the basal stage of the hyperinsulinemic-euglycemic clamp procedure from the periumbilical area by aspiration through a 3-mm liposuction cannula (Tulip Medical Products) connected to a 30 cc syringe. Samples were immediately rinsed in ice-cold saline and frozen in liquid nitrogen before being stored at -80°C until final analyses.

### Hepatic de novo lipogenesis

Subjects consumed 50-mL aliquots of 70% D_2_O (Sigma-Aldrich), provided in sterile vials, every day for 3 to 5 weeks. Aliquots of D_2_O were consumed 3-4 times/day every day for the first 5 days (priming period) followed by two 50-mL doses daily. A blood sample was obtained after an overnight fast the day after the final dose of D_2_O was consumed to determine body water D_2_O enrichment and to measure hepatic DNL by gas chromatography/ mass spectrometry (GC- MS)^34,46^. Compliance with D_2_O consumption was monitored by interview at weekly visits with the study research coordinator, by counting the return of empty vials at each visit, and by evaluating D_2_O enrichments in plasma (obtained on day 7 and weekly thereafter) and saliva (obtained on days 2, 4, and 11 and then weekly thereafter).

### Adipose tissue RNA sequencing

Total RNA was isolated from frozen adipose tissue samples by using QIAzol and an RNeasy mini kit in combination with an RNase-free DNase Set (Qiagen)^47^. Library preparation of the samples was performed with mRNA reverse transcribed to yield cDNA fragments, which were then sequenced by using an Illumina NovaSeq 6000 at the UC San Diego IGM Genomics Center. Expression of *LPIN1* is presented as log_2_-transformed counts per million reads. The RNA sequencing data have been deposited in the Gene Expression Omnibus database, https://www.ncbi.nlm.nih.gov/geo (accession no. GSE156906).

### Sample analysis and calculations

Plasma glucose concentration was determined by using an automated glucose analyzer. Plasma insulin, HbA1c, and lipid profile were measured in the Washington University Core Laboratory for Clinical Studies. Deuterium enrichment in total body water, deuterium enrichment and labeling pattern in TG-palmitate, and [U-^13^C]glucose enrichment in plasma glucose were determined by using GC-MS as described previously^48,49^. The hepatic insulin sensitivity index (HISI) was calculated as the inverse of the product of plasma insulin concentration and the endogenous glucose rate of appearance (Ra) in the systemic circulation, determined by dividing the glucose tracer infusion rate by the average plasma glucose tracer-to-tracee ratio (TTR) during the last 20 minutes of the basal period of the HECP^31^. The total glucose rate of disappearance (Rd) during insulin infusion was assumed to be equal to the sum of the endogenous glucose Ra and the rate of infused glucose during the last 20 minutes of the HECP^31^. An index of whole-body insulin sensitivity was calculated as glucose Rd per kg FFM divided by plasma insulin during the hyperinsulinemic-euglycemic clamp procedure. The fractional contribution of DNL to palmitate in plasma TG was calculated by mass isotopomer distribution analysis as described previously^46,48^.

### Mouse models

All animal studies were approved by the Institution Animal Care and Use Committee of Washington University. Male and female mice were maintained in the C57BL/6J background. Mice were group housed, given free access to a standard chow diet unless otherwise stated, and maintained on a standard 12 h light-dark cycle. Mice harboring a *Lpin1* floxed allele were previously described (B6(Cg)-*Lpin1^tm1c(EUCOMM)HMGU^*/FincJ; available from Jackson Laboratory, strain number 032117)^27^. Adipocyte-specific knockout mice were generated by crossing *Lpin1* floxed mice with mice expressing adiponectin promoter-driven Cre recombinase (B6;FVB-Tg(Adipoq-cre)1Evdr/J Jackson Laboratory, strain number 028020). Littermate control mice were homozygous for *Lpin1* floxed alleles but did not express Cre recombinase.

Prior to sacrifice, all mice were fasted for 4 hours starting at 0900. Mice were euthanized via CO_2_ asphyxiation. Blood was procured from the inferior vena cave into EDTA coated tubes and plasma was separated via centrifugation. All tissues were excised and immediately flash-frozen in liquid nitrogen, and stored at -80°C until further use.

### Diet studies

Male mice were given *ad libitum* access to the indicated diets at 8-weeks of age. A high-fat diet (HFD, 60 kcal %, Research Diets, D12492) or a matched sucrose low-fat diet (LFD, 10 kcal %, Research Diets, D12450J) was provided for 5 weeks. For the NASH studies, a diet high in fructose (17 kcal %), fat (palm oil 40 kcal %), and cholesterol (2%) (HFHF-C, Research Diets, D09100310) diet or a matched sucrose, high-sugar (dextrose), low-fat (10 kcal % (HSLF, Research Diets, D09100304) diet was provided for 16-weeks.

### Glucose and insulin tolerance tests

Prior to all tolerance tests, mice were placed on hardwood bedding and fasting was initiated at 0900 h. Glucose tolerance tests (GTT) were performed in 5-hour fasted mice. Glucose was dissolved in saline and given via an intraperitoneal injection (1 g glucose/kg lean mass). Insulin tolerance tests (ITT) were performed in 4-hour fasted mice. Recombinant human insulin (Humalin R, Eli Lilly; 0.75 U/kg lean mass) in saline was given via an intraperitoneal injection. Blood was procured from the tail and glucose was monitored via a glucometer (One Touch Ultra, Life Scan Inc.) at times indicated. Lean and fat mass were determined by EchoMRI prior to tolerance testing to determine lean mass for injections.

### Mouse hyperinsulinemic-euglycemic clamp studies

Hyperinsulinemic-euglycemic clamp studies were performed at the Vanderbilt Mouse Metabolic Phenotyping Center. All procedures required for the hyperinsulinemic–euglycemic clamp were approved by the Vanderbilt University Animal Care and Use Committee. Catheters were implanted into a carotid artery and a jugular vein of mice for sampling and infusions respectively five days before the study as described by Berglund et al ^50^. Insulin clamps were performed on mice fasted for 5 hours using a modification of the method described by Ayala et al ^51^. [3-^3^H]-glucose was primed (1.5μCi) and continuously infused for a 90 min equilibration and basal sampling periods (0.075 μCi/min). [3-^3^H]-glucose was mixed with the non-radioactive glucose infusate (infusate specific activity of 0.5 μCi/mg) during the 2-hour clamp period. Arterial glucose was clamped using a variable rate of glucose (plus trace [3-^3^H]-glucose) infusion, which was adjusted based on the measurement of blood glucose at 10 min intervals. By mixing radioactive glucose with the non-radioactive glucose infused during a clamp, deviations in arterial glucose specific activity are minimized and steady state conditions are achieved. The calculation of glucose kinetics is therefore more robust ^52^. Baseline blood or plasma variables were calculated as the mean of values obtained in blood samples collected at −15 and −5 min. At time zero, insulin infusion (4 mU/kg body weight/min) was started and continued for 120 min. Mice received heparinized saline-washed erythrocytes from donors at 5 μl/min to prevent a fall in hematocrit. Blood was taken from 80–120 min for the determination of [3-^3^H]-glucose. Clamp insulin was determined at *t*=100 and 120 min. At 120 min, 13µCi of 2[^14^C]deoxyglucose ([^14^C]2DG) was administered as an intravenous bolus. Blood was taken from 2-25 min for determination of [^14^C]2DG. After the last sample, mice were anesthetized and tissues were freeze-clamped for further analysis. Plasma insulin was determined by radioimmunoassay at 10 minutes prior to clamp (Fasting) and averaged at 90 and 120 minutes after the start of the clamp (Clamp). Radioactivity of [3-^3^H]-glucose and [^14^C]2DG in plasma samples, and [^14^C]2DG-6-phosphate in tissue samples were determined by liquid scintillation counting. Glucose appearance (Ra) and disappearance (Rd) rates were determined using steady-state equations^53^. Endogenous glucose appearance (endoRa) was determined by subtracting the GIR from total Ra. The glucose metabolic index (Rg) was calculated as previously described ^54^.

### Metabolic cage studies

Metabolic measurements were conducted as previously described ^55^. Eight-week old male mice were fed a LFD (*n* = 5-7) or HFD (*n* = 5-7) for 5 weeks prior to body composition analysis via EchoMRI and single-housing in a 16-metabolic cage Comprehensive Laboratory Animal Monitoring System (CLAMS, Columbus Instruments). Mice were acclimated overnight prior to measurements starting at 0600 h to 0559 h for 48 hours (Zeitgeber time 0hr to 48hr). Mice were housed at room temperature on a standard 12 hour light/dark cycle and staggered to equally distribute genotypes across the cages. Mice were given *ab libitum* access to food (LFD or HFD) and water (hanging feeders and water bottles on load cells). Cumulative food intake was determined over a 48 hour period. Activity was monitored as infrared laser array beam breaks on an X- and Y-axis. Oxygen (O_2_) consumption and carbon dioxide (CO_2_) production were measured by indirect calorimetry using a zirconia O2 sensor and CO2 sensor at air flow rates of 0.90 L/min (18 sec line bleed followed by 2 sec measurement for each cage). Cages were measured individually in series (∼5.5 min for each complete interval), with room air sampled between each interval to calculate O_2_ consumption and CO_2_ production. Data were analyzed by the web-based indirect calorimetry analysis tool, CalR^33^ to determine the effect of weight differences on metabolic data.

### Histology and NAFLD scoring

Tissues were harvested as described above. Tissue was fixed in 10% neutral buffered formalin for 48 hours. Samples were rinsed and stored in 70% EtOH until paraffin embedding, sectioning, and staining with hematoxylin-eosin (H&E) in the Anatomic Molecular Pathology Core Labs at Washington University School of Medicine. NAFLD scoring was conducted by a blinded independent clinical pathologist and the NAFLD total score (x/8) was a summation of the steatosis score (x/3), lobular inflammation (x/3), and ballooning (x/2)^56^.

### Immunoblotting

Thirty micrograms of protein were loaded onto 4-15% acrylamide pre-cast gels (BioRad, 64329760) in Lamellae loading buffer (BioRad, 161-0737). Proteins were transferred to PVDF membranes in Tris-glycine buffer with 10% methanol. Following transfer, membranes were blocked in 5% BSA in TBS for 1 hour prior to overnight antibody incubations in 5% BSA in TBS. Antibodies used are as follows: lipin 1 (rabbit, Santa Cruz Biotechnology, sc-98450), lipin 2 (rabbit, as previously described^57^), and β-actin (mouse, Sigma Aldrich, SAB4502543). Following incubations, membranes were washed and incubated in secondary antibodies (LiCor) prior to imaging on a LiCor Odyssey.

### Liver triglyceride and plasma analytes

Flash frozen liver tissue was thawed and homogenized in ice cold PBS (100 mg/ mL). Lipids were solubilized in 1% sodium deoxycholate via vortexing and heating at 37°C for 5 minutes. Triglycerides were determined enzymatically using the Infinity triglyceride colorimetric assay according to the manufacturer’s instructions (Thermo Fisher, TR22421). Blood from 4 hour fasted mice was collected into ethylenediaminetetraacetic acid (EDTA) coated tubes. Plasma was separated via centrifugation at 8,000 x g for 8 minutes at 4°C. Plasma lipids were determined using commercially available colorimetric assays as follow: triglycerides (Thermo Fisher, TR22421), Non-esterified free fatty-acids (NEFAs, WAKO, XYZ), and free glycerol (Sigma, F26248). Plasma alanine transaminase (ALT) and aspartate aminotransferase (AST) were measured using liquid kinetic assays (TECO Diagnostic, A534 and A559). Plasma insulin was determined by Singlex Immunoassay at the Washington University Core Laboratory for Clinical Studies. Plasma adiponectin and adipokines (leptin, resistin, and TNFα) were determined by the Cellular Molecular Biology Core at the Washington University Nutrition Obesity Research Center using suspension magnetic bead based Singlex (adiponectin) or Multiplexed Immunoassays (MilliporeSigma, MADPNMAG-70K-01, and MADKMAG-71K) following manufacturer’s protocol and were analyzed using a Luminex 200 system (Luminex Corp.)

### *De novo* hepatic lipogenesis

Total liver palmitate was analyzed for mass isotope abundances by LC-MS, using MIDA to determine the effective body water deuterium exposure (precursor pool enrichment) for the calculation of fractional DNL^34,58,59^. Briefly, animals from each group were labeled with an intraperitoneal injection of 100% ^2^H_2_O saline (35 µl/g body weight ∼3.5 % of body weight, deuterium oxide, Sigma 151890) and provided with 8% ^2^H_2_O drinking water for the remainder of the study to maintain body ^2^H_2_O enrichments of approximately 5%. Livers were then homogenized in acetonitrile, methanol and water (2:2:1) for fatty acid solubilization and subsequent analysis by LC-MS to quantify the isotopic enrichment (M1, M2, M3, M4) due to the incorporation of deuterium from heavy water into palmitate. Specifically, a 2 μl aliquot containing the fatty acid metabolites was subjected to LC/MS analysis by using an Agilent 1290 Infinity II (LC) system coupled to an Agilent 6545 Quadrupole-Time-of-Flight (QTOF) mass spectrometer with a dual Agilent Jet Stream electrospray ionization source. Samples were separated on a SeQuant ZIC-pHILIC column (100 3 2.1 mm, 5 mm, polymer, Merck-Millipore) including a ZIC-pHILIC guard column (2.1 mm x 20 mm, 5 mm) To confirm only palmitate eluted at its retention time, samples were analyzed with higher resolution (120,000) on an Agilent ID-X Orbitrap. The column compartment temperature was maintained at 40C and the flow rate was set to 250 mL/min. The mobile phases consisted of A: 95% water, 5% acetonitrile, 20 mM ammonium bicarbonate, 0.1% ammonium hydroxide solution (25% ammonia in water), 2.5 mM medronic acid, and B: 95% acetonitrile, 5% water, 2.5 mM medronic acid. The following linear gradient was applied: 0 to 1 min, 90% B; 12 min, 35% B; 12.5 to 14.5 min, 25% B; 15 min, 90% B followed by a re-equilibration phase of 4 min at 400 mL/min and 2 min at 250 mL/min. Palmitate ions were monitored at a mass of 256.240, 257.244, 258.247, and 259.248, representing the parent M0 through M3 isotopes. Excess M1 and M2 enrichments were determined by subtraction of the natural abundance values in unlabeled standards (run in parallel). The proportion of plasma palmitate that originated from the DNL pathway was then calculated from the excess M1 and M2 of palmitate by using MIDA to determine both the biosynthetic precursor enrichment and the corresponding isotopic enrichment of newly synthesized palmitate molecules^59^. The precursor pool enrichment (p) was determined from the ratio of EM2/EM1 in the experimental data. Knowledge of the calculated metabolic precursor pool enrichment and the known n (number of repeating subunits in the polymer = 21 for palmitate^60^ allowed calculation of the theoretical asymptote enrichment of the single-labeled mass isotopomer species (excess M1), representing the maximum possible enrichment when palmitate is newly synthesized at this deuterium precursor pool enrichment.

### Lipidomic analysis

Tissue and plasma lipidomic analysis was performed at the Washington University Metabolomics Facility as previously described ^61^. The liver and gastrocnemius muscle samples were homogenized in water (4 mL/g tissue). All the lipids were extracted from 50 µL of plasma or homogenate using protein precipitation method. The DAG, and TAG were further extracted with modified Bligh-Dyer method. DAG (21:0-21:0) (2 µg/sample), and TAG (17:1-17:1-17:1) (13 µg/sample were used as internal standards n plasma. DAG(21:0-21:0) (0.3 µg/sample) and TAG (17:1-17:1-17:1) were used as internal standards in liver and gastric samples. Internal standards were added to the samples before extraction. Quality control (QC) samples were prepared by pooling the aliquots of the study samples and were used to monitor the instrument stability. The QC was injected between every 5 study samples. Measurement of DAG was performed with a Shimadzu 20AD UFLC system coupled to an AB Sciex API4000 mass spectrometer operated in positive MRM mode. Measurement of TAG was performed with a Shimadzu 20AD UFLC system coupled to an AB Sciex 4000QTRAP mass spectrometer operated in positive multiple reaction monitoring (MRM) mode. Data processing was conducted with Analyst 1.6.3. The relative quantification of lipids was provided, and the data were reported as the peak area ratios of the analytes to the corresponding internal standards. The lipid species showed CV < 15% in QC sample injections.

### mRNA isolation and quantification

For mouse studies, total RNA was isolated from frozen tissue using TRizol Plus RNA purification Kits (Thermo Fischer, 12183555) according to the manufacturer’s instructions. Second, 2 μg of RNA was reverse transcribed into complimentary DNA (cDNA) using Taqman High Capacity reverse transcriptase (Life Technologies, 43038228). Quantitative polymerase chain reaction (qPCR) was performed using Power Sybr Green (Applied Biosystems, 4367659) and measured using an ABI QuantStudio 3 sequence detection system (Applied Biosystems). Results were quantified using the 2^-ΔΔCt^ and shown as relative expression from the control groups. Primer sequences are listed in Extended Data Table 2.

### Bulk RNA sequencing and WGCNA analysis

Bulk RNA sequencing was performed at the Genomic Technologies and Access Center at the McDonald Genomic Institute of Washington University School of Medicine in St. Louis. Samples were prepared according to library kit manufacturer’s protocol, indexed, pooled, and sequenced on an Illumina HiSeq. Basecalls and demultiplexing were performed with Illumina’s bcl2fastq software and a custom python demultiplexing program with a maximum of one mismatch in the indexing read. RNA-seq reads were then aligned to the Ensembl release 76 primary assembly with STAR version 2.5.1a ^62^. Gene counts were derived from the number of uniquely aligned unambiguous reads by Subread:featureCount version 1.4.6-p5 ^63^. Isoform expression of known Ensembl transcripts were estimated with Salmon version 0.8.2 ^64^. Sequencing performance was assessed for the total number of aligned reads, total number of uniquely aligned reads, and features detected. The ribosomal fraction, known junction saturation, and read distribution over known gene models were quantified with RSeQC version 2.6.2 ^65^.

All gene counts were then imported into the R/Bioconductor package EdgeR ^66^ and TMM normalization size factors were calculated to adjust for samples for differences in library size. Ribosomal genes and genes not expressed in the smallest group size minus one samples greater than one count per million were excluded from further analysis. The TMM size factors and the matrix of counts were then imported into the R/Bioconductor package Limma ^67^. Weighted likelihoods based on the observed mean-variance relationship of every gene and sample were then calculated for all samples with the voomWithQualityWeights ^68^. The performance of all genes was assessed with plots of the residual standard deviation of every gene to their average log-count with a robustly fitted trend line of the residuals. Differential expression analysis was then performed to analyze for differences between conditions and the results were filtered for only those genes with Benjamini-Hochberg false-discovery rate adjusted *p*-values less than or equal to 0.05.

The heatmap was generated using iDEP 9.0 ^69^. For each contrast extracted with Limma, global perturbations in known Gene Ontology (GO) terms, MSigDb, and KEGG pathways were detected using the R/Bioconductor package GAGE^70^ to test for changes in expression of the reported log 2 fold-changes reported by Limma in each term versus the background log 2 fold-changes of all genes found outside the respective term. Perturbed KEGG pathways where the observed log 2 fold-changes of genes within the term were significantly perturbed in a single-direction versus background or in any direction compared to other genes within a given term with p-values less than or equal to 0.05 were rendered as annotated KEGG graphs with the R/Bioconductor package Pathview ^71^.

To find the most critical genes, the raw counts were variance stabilized with the R/Bioconductor package DESeq2^72^ and was then analyzed via weighted gene correlation network analysis with the R/Bioconductor package WGCNA ^35^. Briefly, all genes were correlated across each other by Pearson correlations and clustered by expression similarity into unsigned modules using a power threshold empirically determined from the data. An eigengene was then created for each de novo cluster and its expression profile was then correlated across all coefficients of the model matrix. Because these clusters of genes were created by expression profile rather than known functional similarity, the clustered modules were given the names of random colors where grey is the only module that has any pre-existing definition of containing genes that do not cluster well with others. These de-novo clustered genes were then tested for functional enrichment of known GO terms with hypergeometric tests available in the R/Bioconductor package clusterProfiler ^73^. Significant terms with Benjamini-Hochberg adjusted p-values less than 0.05 were then collapsed by similarity into clusterProfiler category network plots to display the most significant terms for each module of hub genes in order to interpolate the function of each significant module. The information for all clustered genes for each module were then combined with their respective statistical significance results from Limma to determine whether or not those features were also found to be significantly differentially expressed. Lipidomic and phenotypic data were tested for associations with WGCNA pathways using Pearson’s Correlation Coefficient and significant correlations were considered if *p* < 0.05.

### Statistics

Data are reported as means ± SEM unless otherwise noted. A *p*-value ≤0.05 was considered statistically significant.

For studies conducted in human subjects, statistical analyses were performed by using SPSS (version 28). One-way analysis of variance was used to compare subject characteristics and adipose tissue *LPIN1* expression among MHL, MHO and MUO groups with Fisher’s least significant difference post-hoc procedure used to identify significant mean differences where appropriate. Relationships between adipose tissue *LPIN1* expression and metabolic variables were evaluated by using linear and nonlinear regression analysis with the best-fit to the data reported.

Data obtained from studies conducted in mice was analyzed by using GraphPad Prism software. Independent and paired Student’s T test, Two-Way Analysis of Variance (ANOVA), were performed where appropriate. Secondary post-hoc analysis found differences in groups using either Tukey’s or Sidak’s multiple comparisons where appropriate. Comparisons and replicate numbers are listed in each figure legends.

## Acknowledgements

Plasma and adipocyte analyses were conducted by the Cellular Molecular Biology Core at the Washington University Nutrition Obesity Research Center P30 DK056341. The lipid analysis was supported by the Washington University Diabetes Research Center (P30 DK020579). Hyperinsulinemic-euglycemic clamps were performed by the Vanderbilt Mouse Metabolic Phenotyping Center (DK059637). The Vanderbilt Hormone Assay and Analytical Core performed the insulin analysis (DK059637 and DK020593).

---

**Extended Data Fig. 1:**
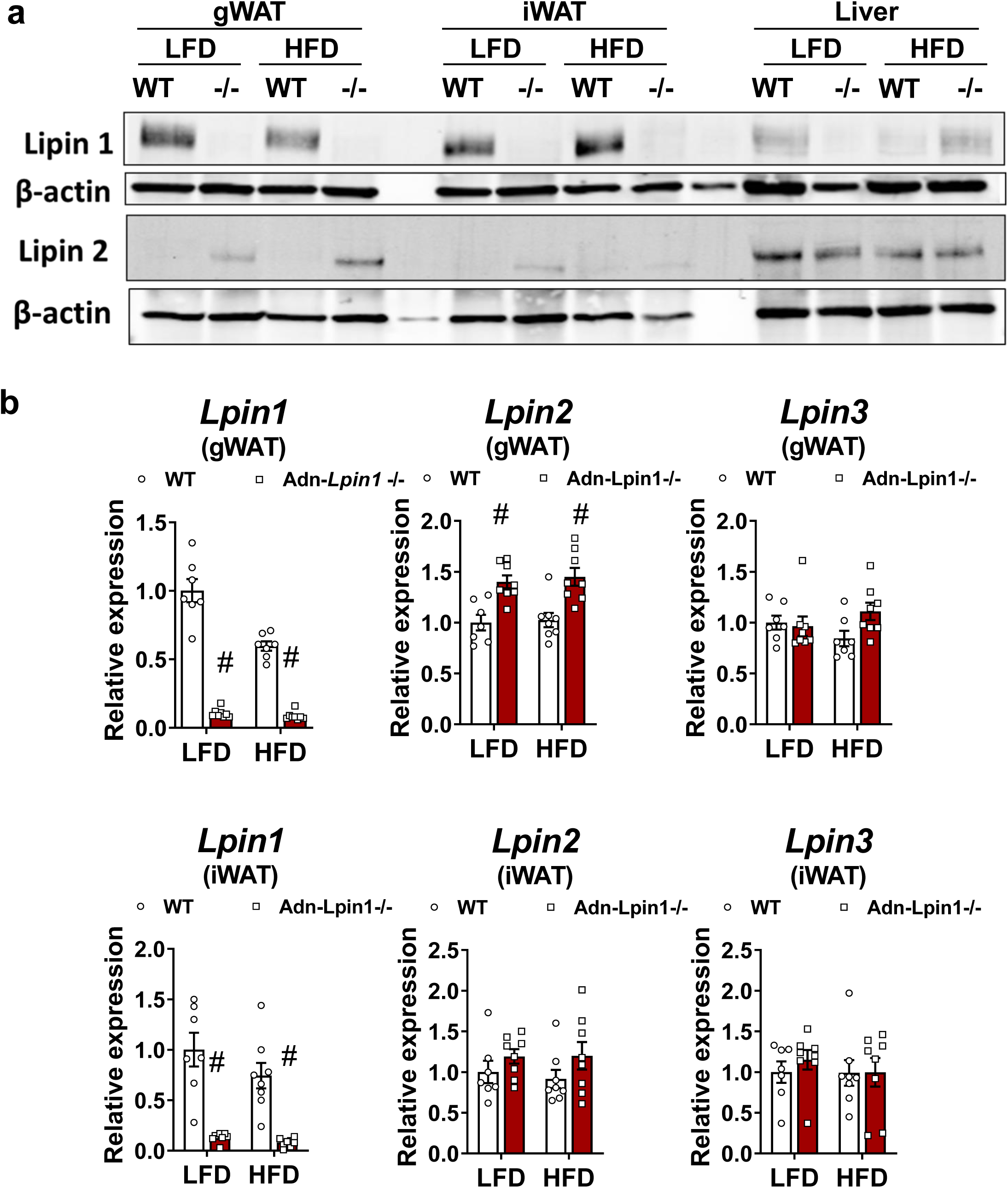
Adn-*Lpin1-/-* mice have a complete loss of lipin 1 in adipose tissue. Eight-week-old male Adn-*Lpin1-/-* mice and their littermate controls (WT) were fed either a 10% low-fat diet (LFD) or a 60% high-fat diet (HFD) for 5 weeks. Mice were fasted for 4 hours prior to sacrifice and tissue collection. **a,** Gonadal white adipose tissue (gWAT), inguinal white adipose tissue (iWAT), and liver tissue proteins were separated via electrophoresis and immunoblotted for lipin 1, lipin 2, and the loading control β-actin. **b,** Gene expression was determined by qPCR and are expressed as relative abundance; lipin 1, lipin 2, and lipin 3 (*Lpin1, Lpin2, Lpin3).* Data are expressed as means +/- S.E.M, and significance was determined by Two-way ANOVA with post-hoc Tukey’s multiple comparisons tests. ^#^*p* < 0.05 for WT vs. Adn-*Lpin1-/-* and ^†^*p* < 0.05 for LFD vs. HFD; (*n* = 7-8).

**Extended Data Fig. 2:**
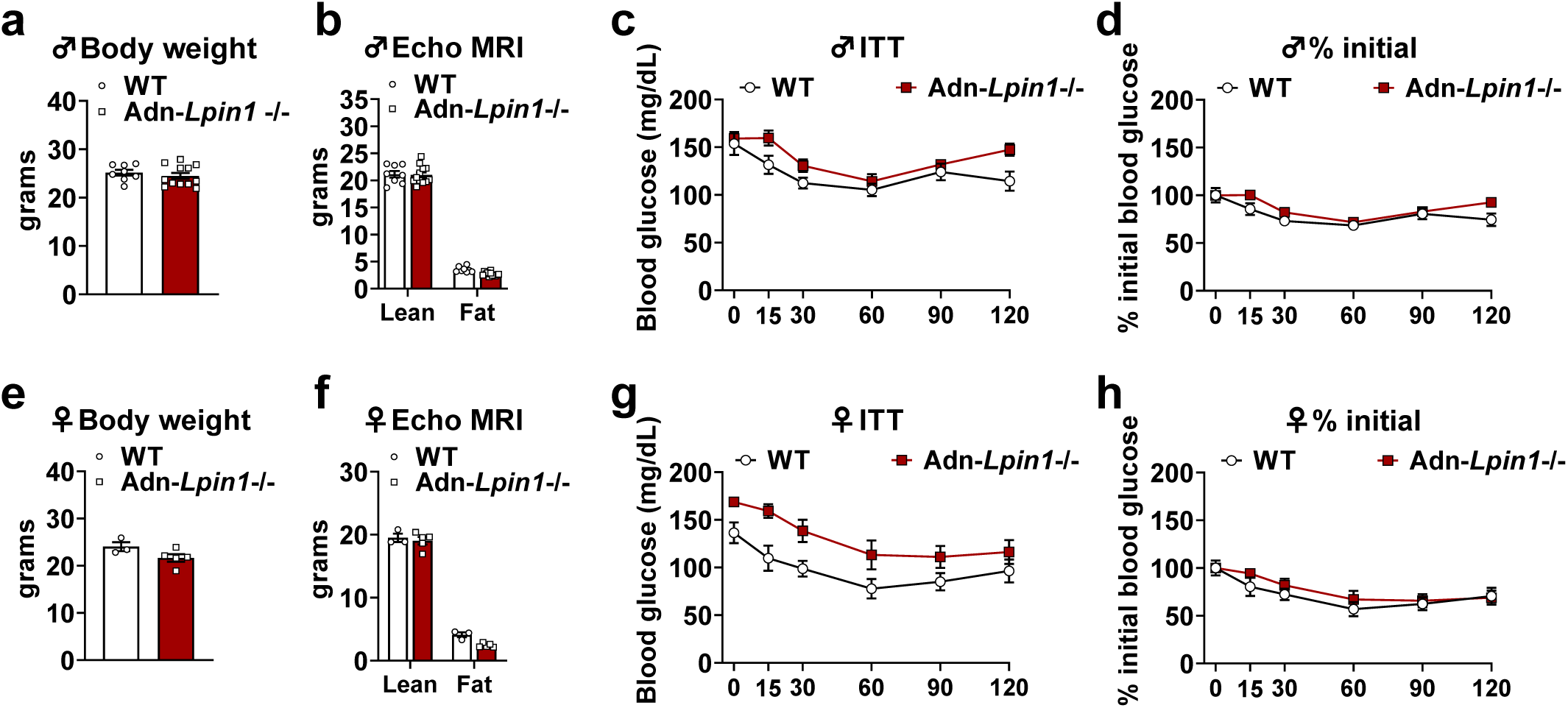
Adn-*Lpin1-/-* mice are outwardly normal on a chow diet. Twelve-week-old male and female Adn-*Lpin1-/-* mice and their wild-type littermate controls (WT) were given ab libitum access to a standard chow diet. **a,** Body weight of fed-male mice. **b,** Male body weight was determined by ECHO MRI. **c,d,** Insulin tolerance tests (ITT) were performed in male mice fasted for 4 hours prior to an intraperitoneal (IP) injection of recombinant human insulin (0.75 U / kg lean mass) and blood glucose was monitored from tail blood at the times indicated. (males *n* = 9-12). **e,h,** Female mice were examined using the same procedures as above. (females *n* = 3-5). Data are expressed as means +/- S.E.M.

**Extended Data Fig. 3:**
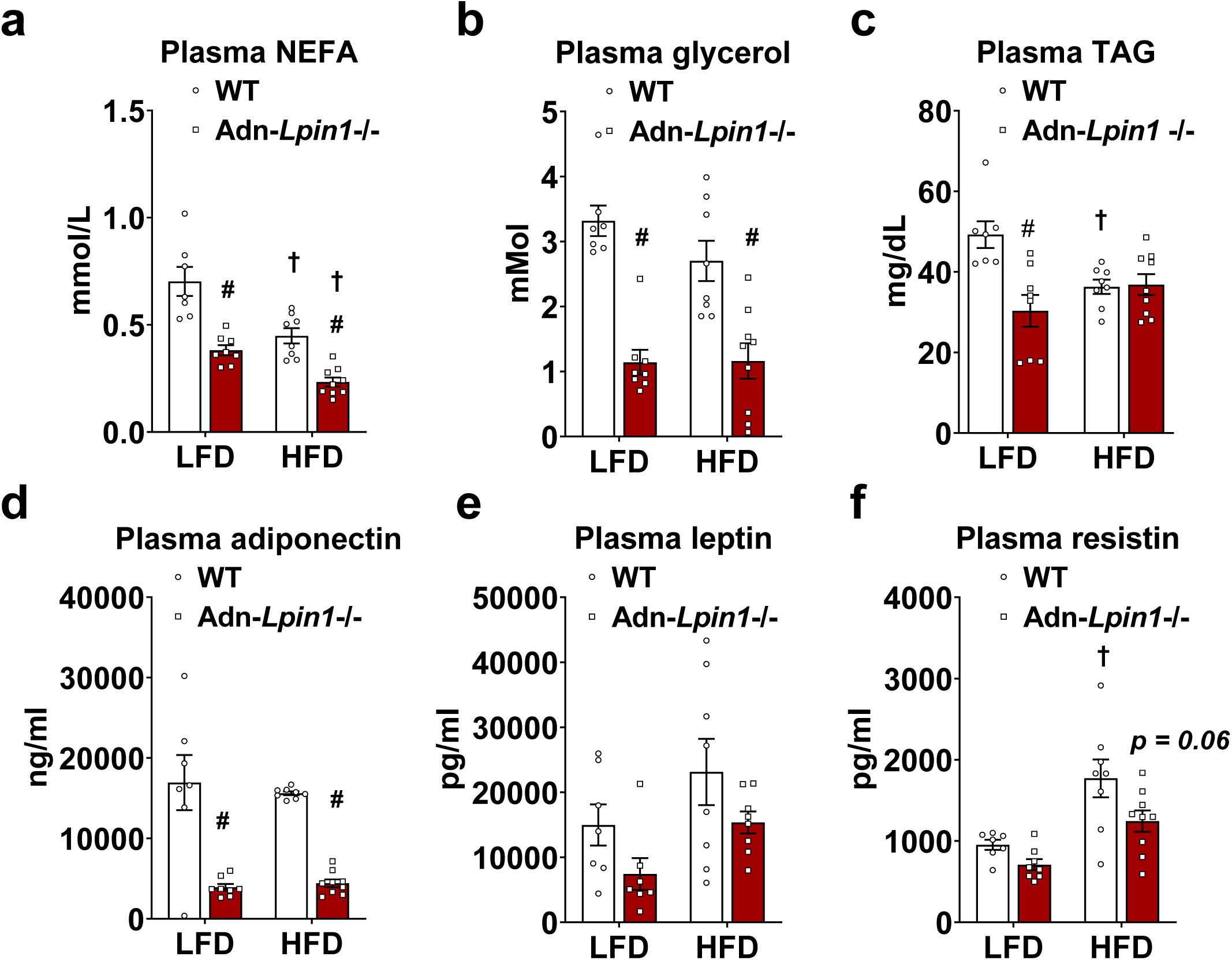
Loss of adipocyte *Lpin1* reduces plasma NEFA and adipokine concentrations. Eight-week-old male Adn-*Lpin1-/-* and control mice were fed either a 10% LFD or a 60% HFD for 5 weeks. Mice were fasted for 4 hours prior to sacrifice and blood was collected into EDTA-coated tubes and plasma was separated via centrifugation. **a-c,** Plasma non-esterified fatty acids (NEFA), glycerol, and triglycerides (TAG) were measured using colorimetric assays according to the manufacturers’ instructions. **d,** Plasma adiponectin was measured using a Singlex Immunoassay. **e,f** Plasma leptin and resistin were measured by Multiplex Immunoassays. Data are expressed as means +/- S.E.M, and significance was determined by Two-way ANOVA with post-hoc Tukey’s multiple comparisons tests. ^#^*p* < 0.05 for WT vs. Adn-*Lpin1-/-* and ^†^*p* < 0.05 for LFD vs. HFD; (*n* = 7-9).

**Extended Data Fig. 4:**
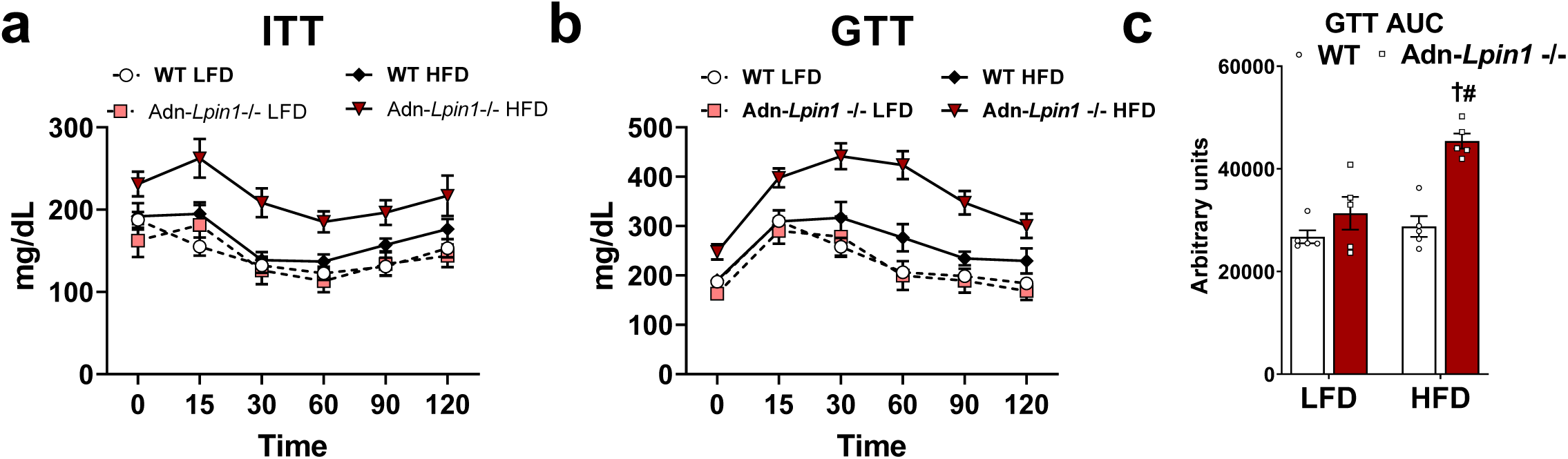
Short-term HFD feeding causes insulin and glucose intolerance in Adn-*Lpin1-/-* mice. Eight-week-old male Adn-*Lpin1-/-* mice and their WT littermate controls were fed either a 10% low-fat diet (LFD) or a 60% high-fat diet (HFD) for 5 weeks. **a,** During week 4 of dietary feeding, lean mass was determined via ECHO MRI, and mice were fasted for 4 hours prior to an insulin tolerance test (ITT) via an intraperitoneal (IP) injection of recombinant human insulin (0.75 U / kg lean mass). **b,** After 5 weeks of diet mice were fasted 5 hours prior to a glucose tolerance test (GTT) via an IP injection of glucose (1 g/ kg lean mass, dissolved in saline. Blood glucose was monitored in tail blood at the times indicated. **c,** Area under the curve was calculated for the GTT. Data are expressed as means +/- S.E.M, and significance was determined by Two-way ANOVA with post-hoc Tukey’s multiple comparisons tests. ^#^*p* < 0.05 for WT vs. Adn-*Lpin1-/-* and ^†^*p* < 0.05 for LFD vs. HFD; (*n* = 5).

**Extended Data Fig. 5:**
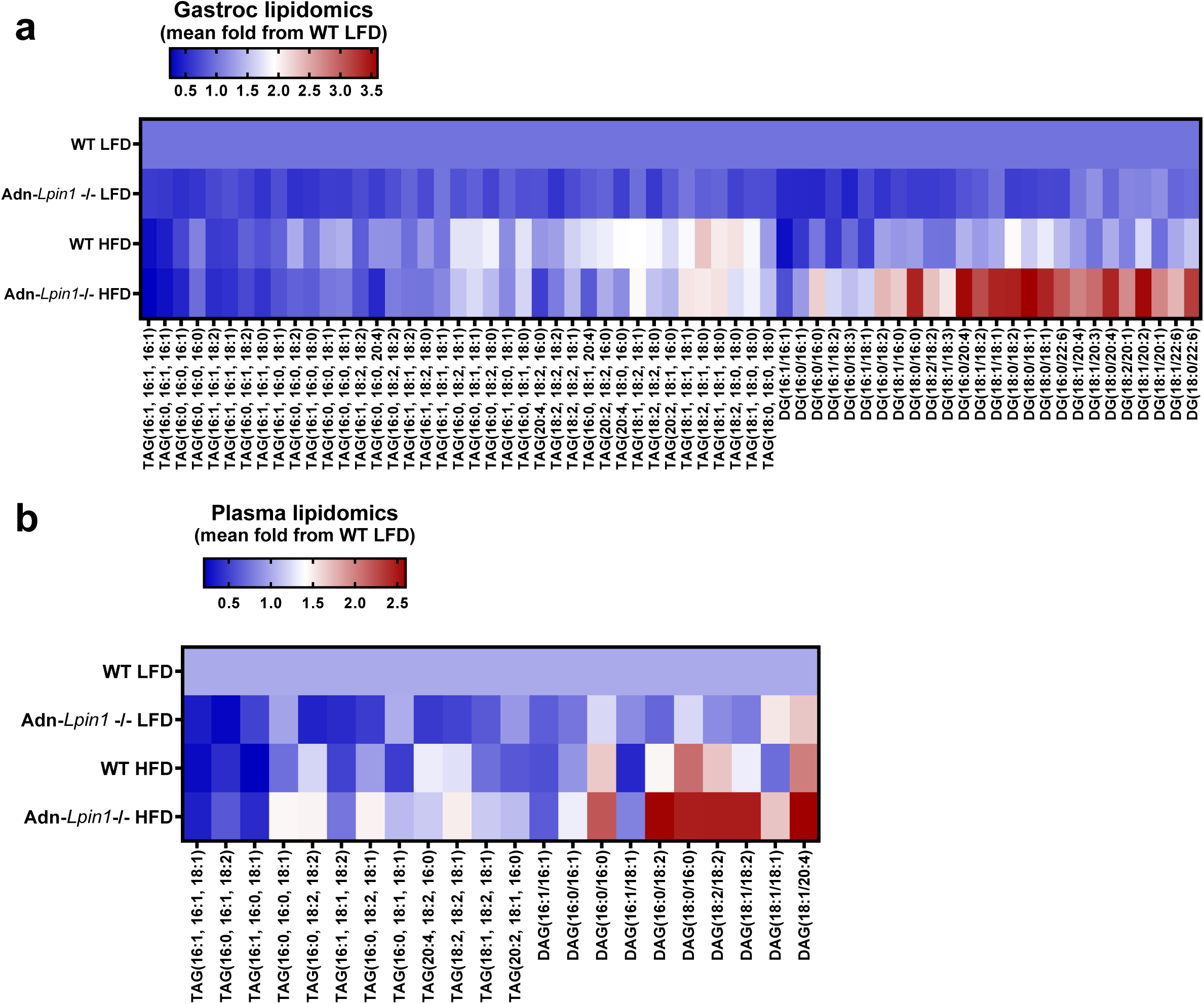
Lipid content of gastrocnemius and plasma. Eight-week-old male Adn-*Lpin1-/-* and WT mice were fed either a 10% LFD or a 60% HFD for 5 weeks. Mice were fasted for 4 hours prior to sacrifice and tissue collection. Triglycerides (TAG) and diacylglycerols (DAG) were extracted and the relative abundance of each species was determined by LC-MS/MS against an internal standard. **a,** Gastrocnemius (gastroc) lipids. **b,** Plasma lipidsData are expressed as mean fold change from the WT LFD-fed mice; (*n* = 5-6).

**Extended Data Fig. 6:**
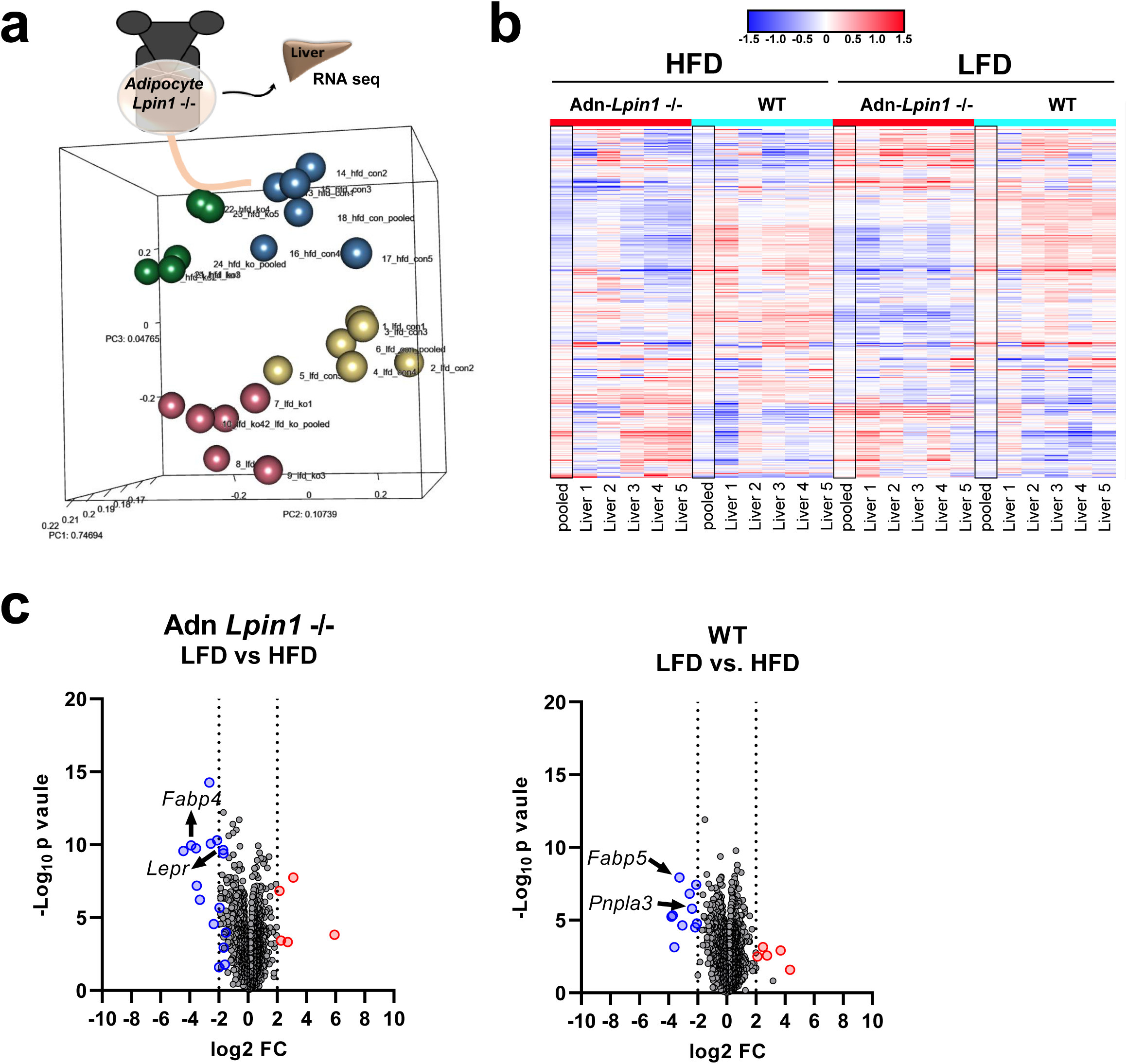
Bulk RNA sequencing in liver. Eight-week-old male Adn-*Lpin1-/-* and control mice were fed either a 10% LFD or a 60% HFD for 5 weeks. Mice were fasted for 4 hours prior to sacrifice, liver collection, RNA isolation, and Bulk RNA sequencing. **a,** PCA plot showing separation of the four groups. **b,** Heatmap of merged differentially expressed data. **c,** Volcano plots of merged differential expression data were graphed as log_2_ fold change versed – log_10_ unadjusted *p-*value. (*n* = 6).

**Extended Data Fig. 7:**
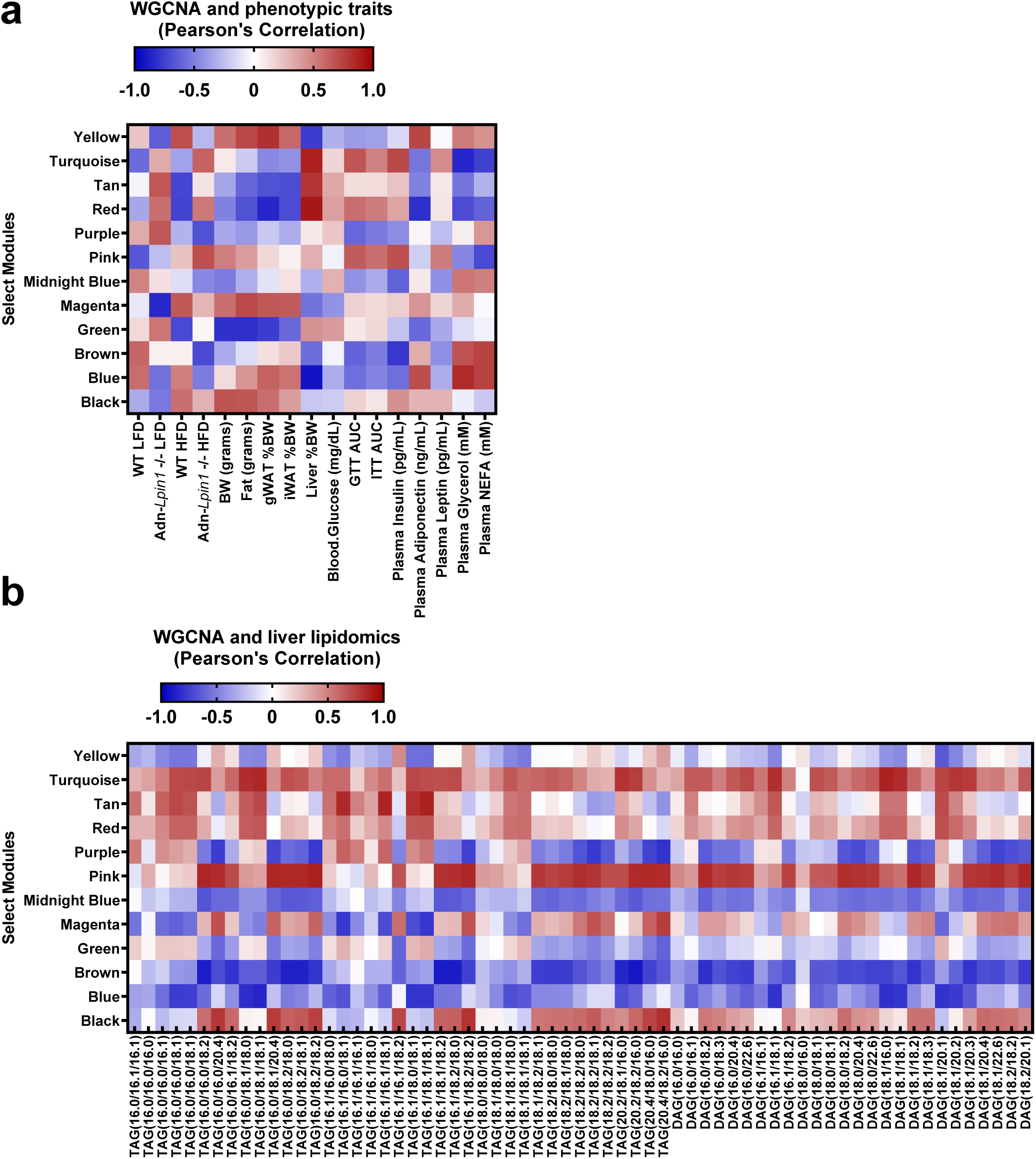
WGCNA module association with and phenotypic data and liver lipidomic data. WGCNA module sets were combined with lipidomic and phenotypic data and associations were tested using Pearson’s Correlation Coefficient; \**p* < 0.05, (*n* = 6). **a,** WGCNA modules and their correlation to phenotypic traits from the mice used to generate the WGCNA data. **b,** WGCNA modules associations with liver lipidomic data.

**Extended Data Fig. 8:**
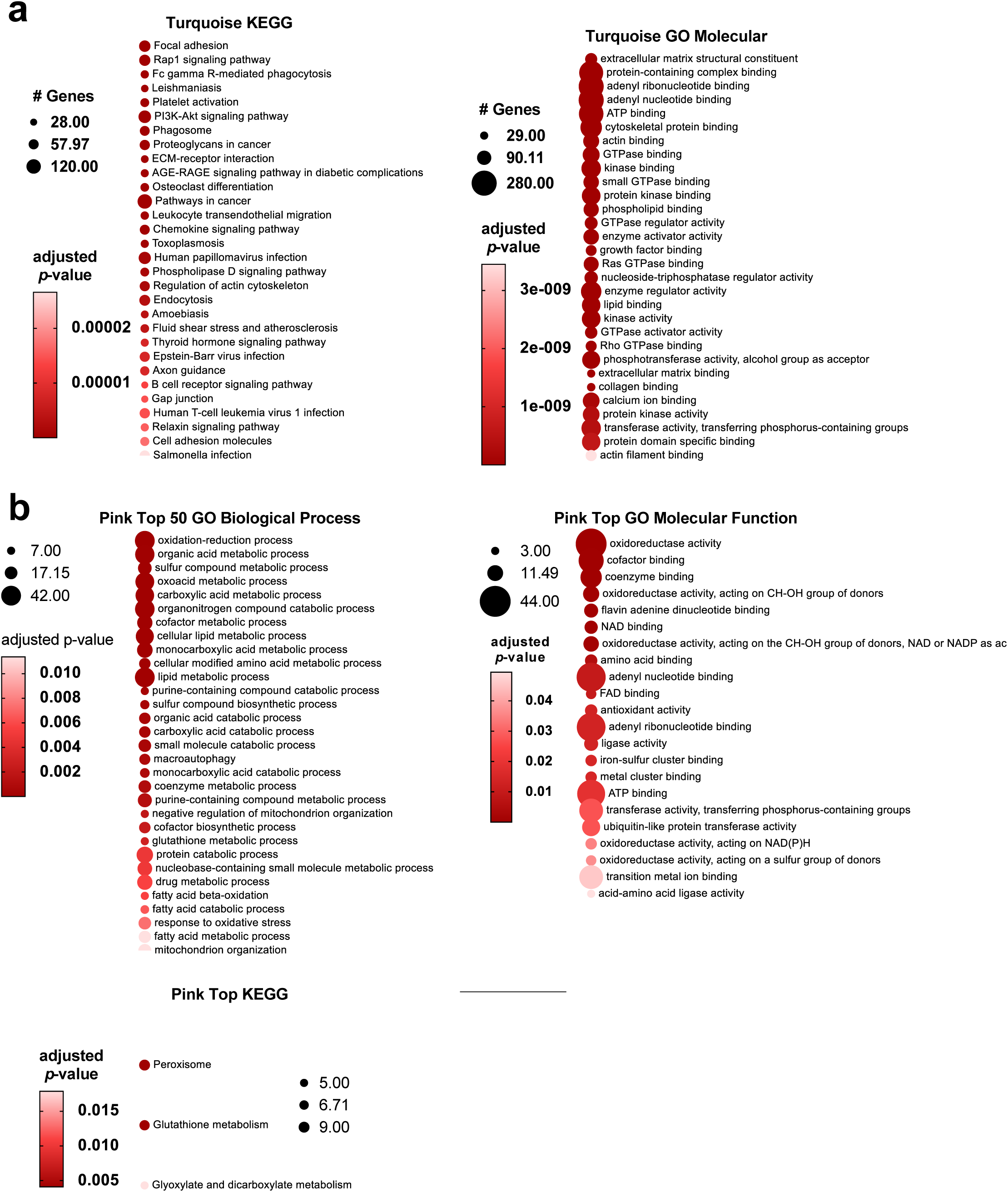
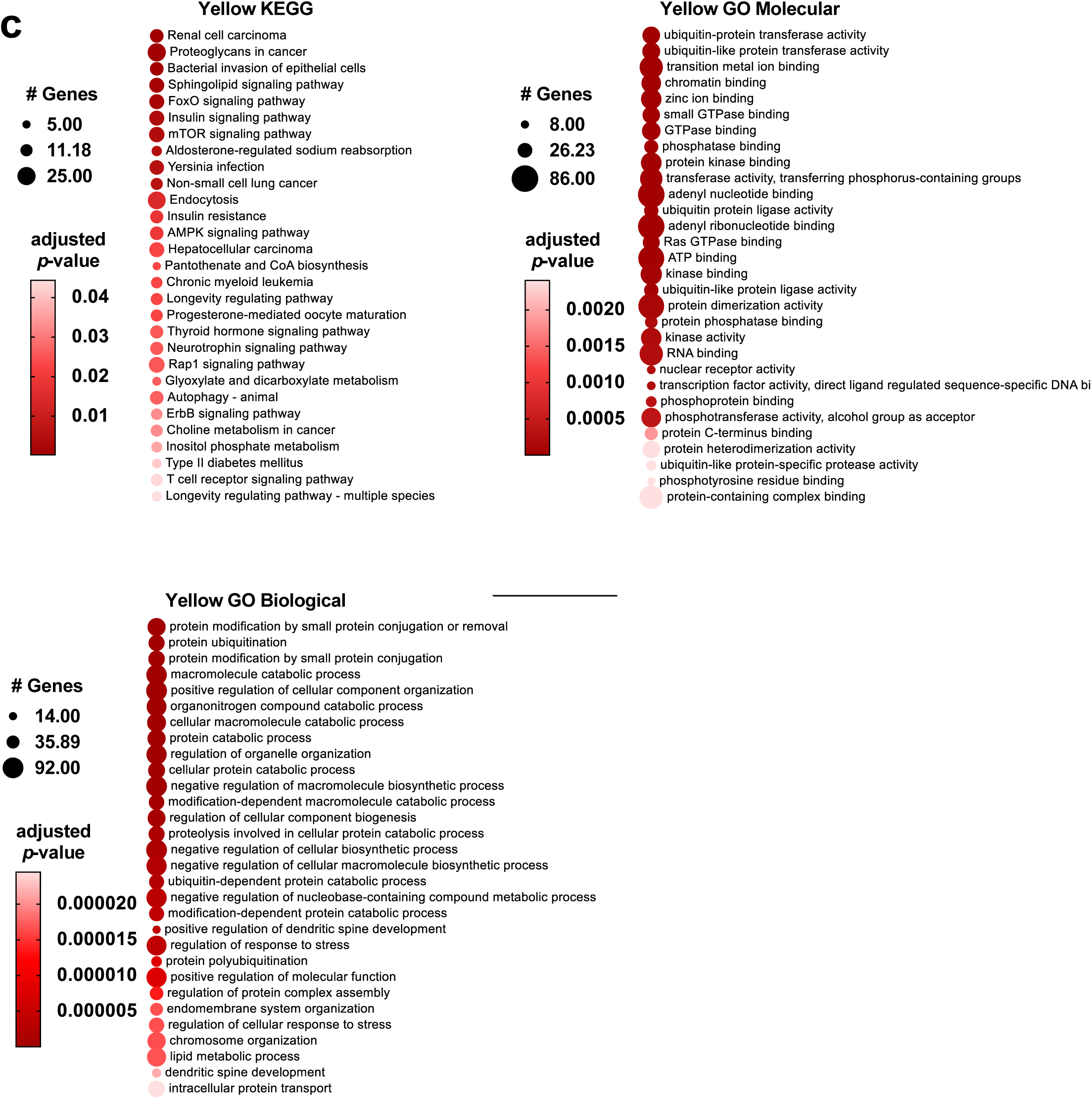
Select WGCNA module set pathway analysis. **a-c,** Graphical representations of top KEGG, and GO pathways in the turquoise, pink, and yellow module sets. The color of the circle represents its adjusted *p-value* and the size represents the number of genes within that pathway.

**Extended Data Fig. 9:**
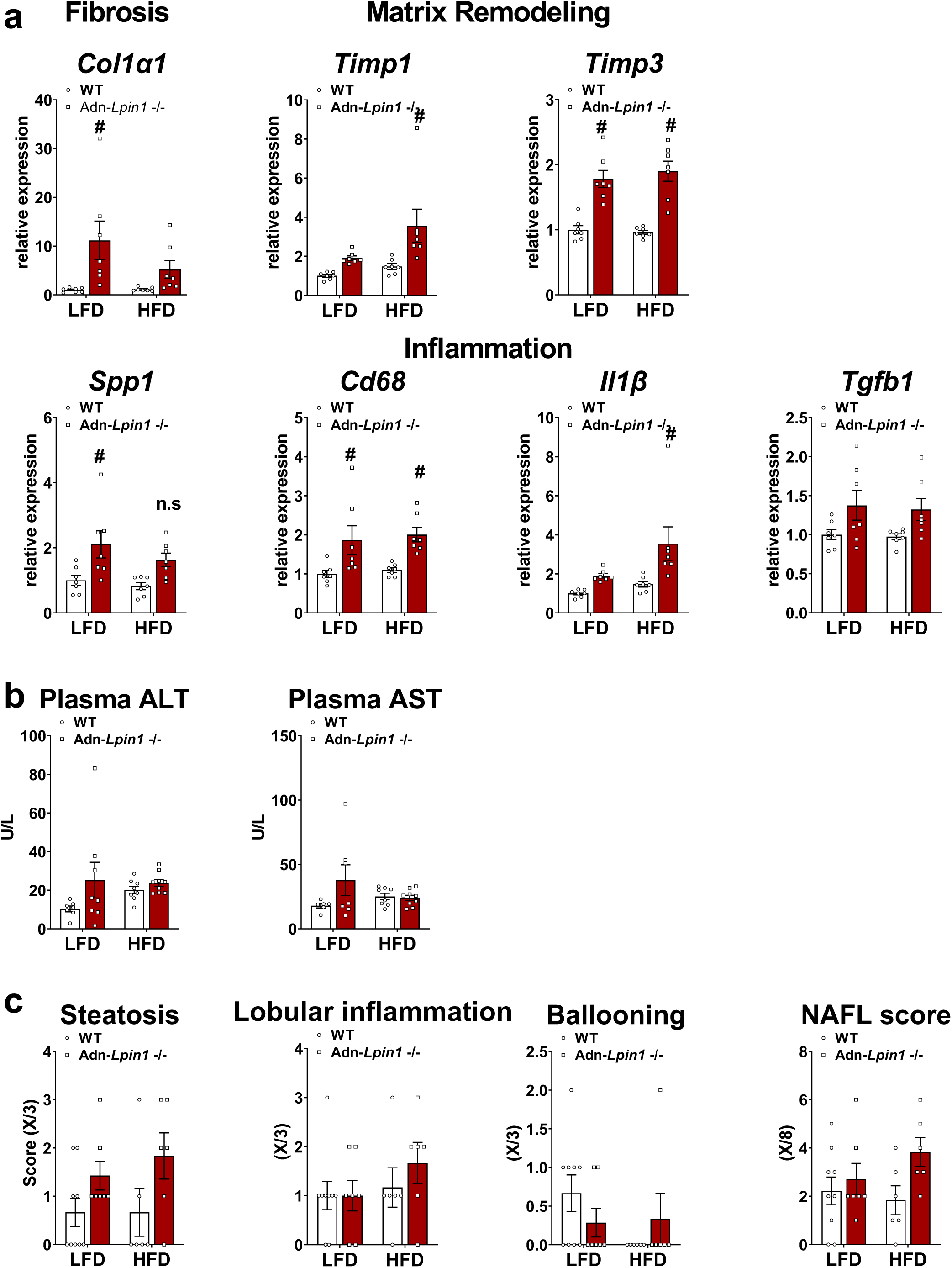
Loss of adipocyte *Lpin1* increases early signs of hepatic fibrosis, matrix remodeling, and inflammation. Eight-week-old male Adn-*Lpin1-/-* and control mice were fed either a 10% LFD or a 60% HFD for 5 weeks. Mice were fasted for 4 hours prior to sacrifice. **a,** Gene expression was determined by qPCR and are expressed as relative abundance; collagen type I alpha 1 chain (*Col1a1*), tissue inhibitor of metalloproteinase 1 & 3 (*Timp1 and Timp3*), secreted phosphoprotein 1 (*Spp1*), cluster of differentiation 68 (*Cd68*), interleukin 1 beta (*Il1b)*, transforming growth factor beta 1 (*TGFB1*). **b,** Plasma alanine transferase (ALT) and aspartate aminotransferase (AST) were measured using liquid kinetic assays. **c,** H&E stained tissue sections were scored by an independent clinical pathologist. Data are expressed as means ± S.E.M., and significance was determined by Two-way ANOVA with post-hoc Tukey’s multiple comparisons tests. ^#^*p* < 0.05 for WT vs. Adn-*Lpin1-/-*; (*n* = 5-9).

**Extended Data Fig. 10:**
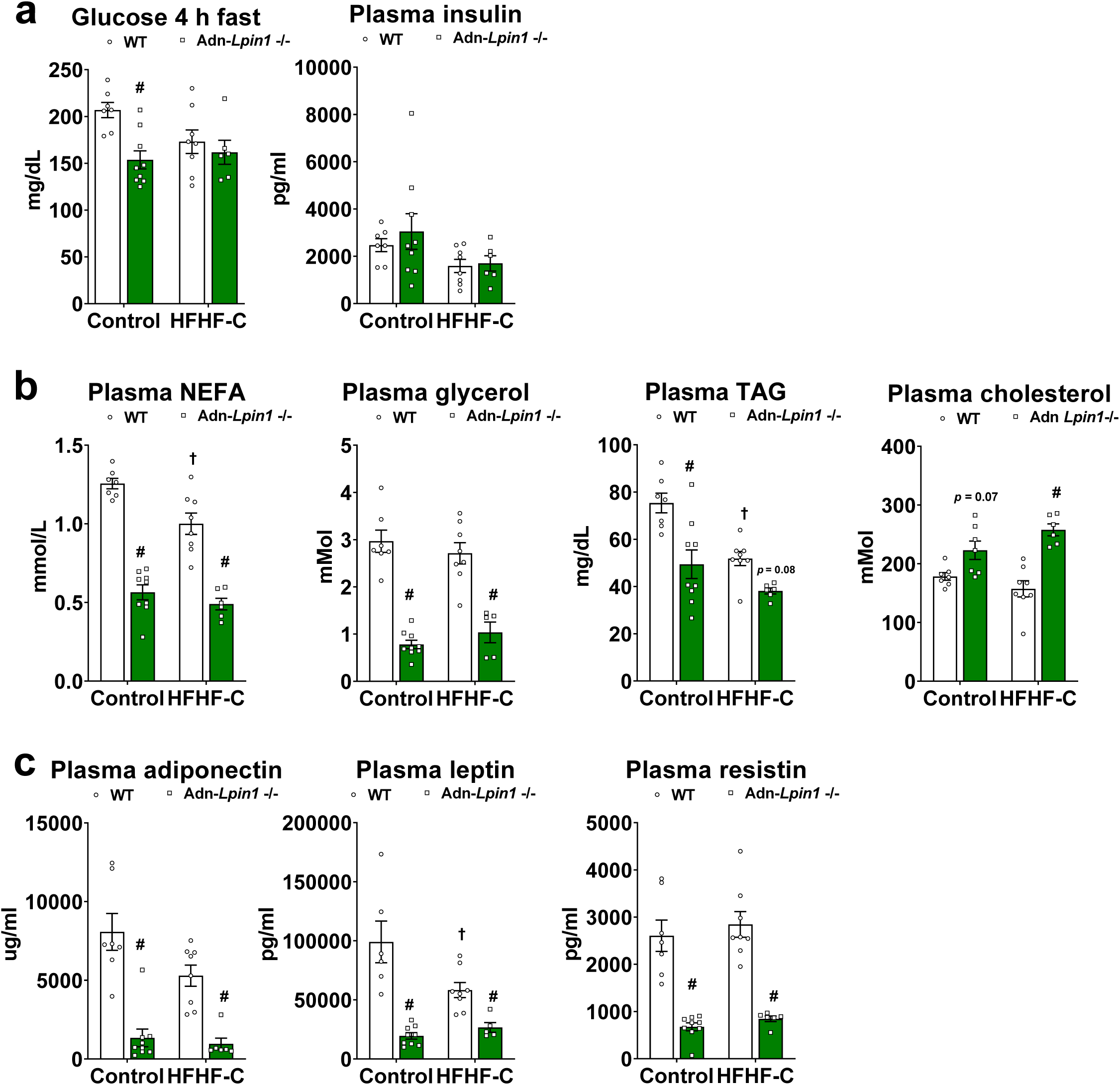
Plasma characteristics of mice fed a NASH-inducing diet. Eight-week-old male mice were fed a diet high in fructose (17 kcal %), fat (palm oil 40 kcal %), and cholesterol (2%) (HFHF-C) or a matched high-sucrose low-fat (10 kcal %) control diet (HSHF) for 16 weeks. Mice were fasted for 4 hours prior to sacrifice and tissue collection. **a,** Blood glucose and plasma insulin concentrations after a 4 hour fast. **b,** Plasma non-esterified fatty acids (NEFA), glycerol, and triglycerides (TAG) were measured using colorimetric assays according to the manufacturers’ instructions. **c,** Plasma adiponectin was measured using a Singlex Immunoassay and plasma leptin and resistin were measured by Multiplex Immunoassays. Data are expressed as means +/- S.E.M, and significance was determined by Two-way ANOVA and post-hoc Tukey’s or Sidak’s multiple comparisons tests. ^#^*p* < 0.05 for WT vs. Adn-*Lpin1-/-* and ^†^*p* < 0.05 for HSLF vs HFHF-C diet; (*n* = 7-9).

**Extended Data Table 1.**
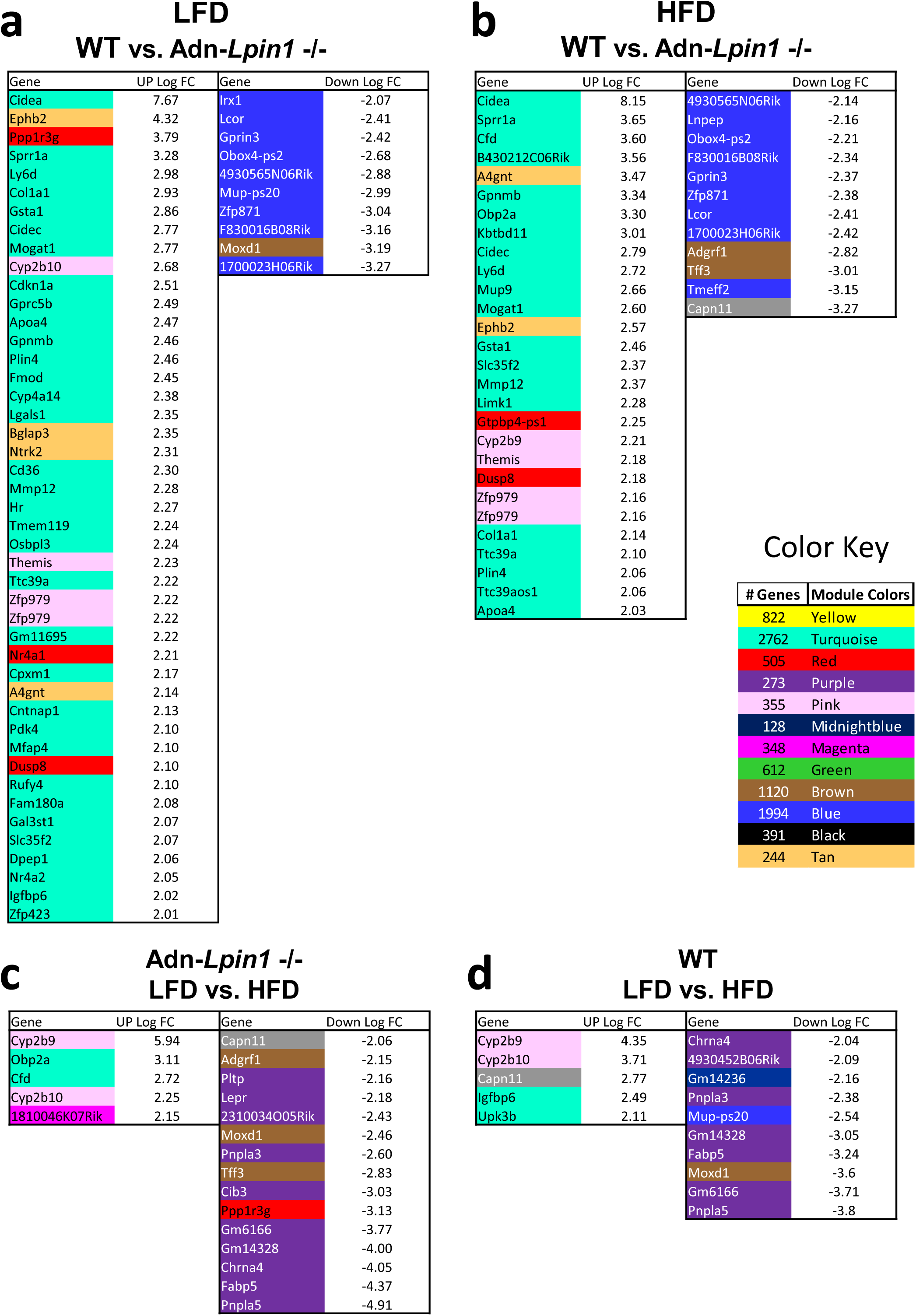
Significantly upregulated genes and their WGCNA module association.

**Extended Data Table 2:**
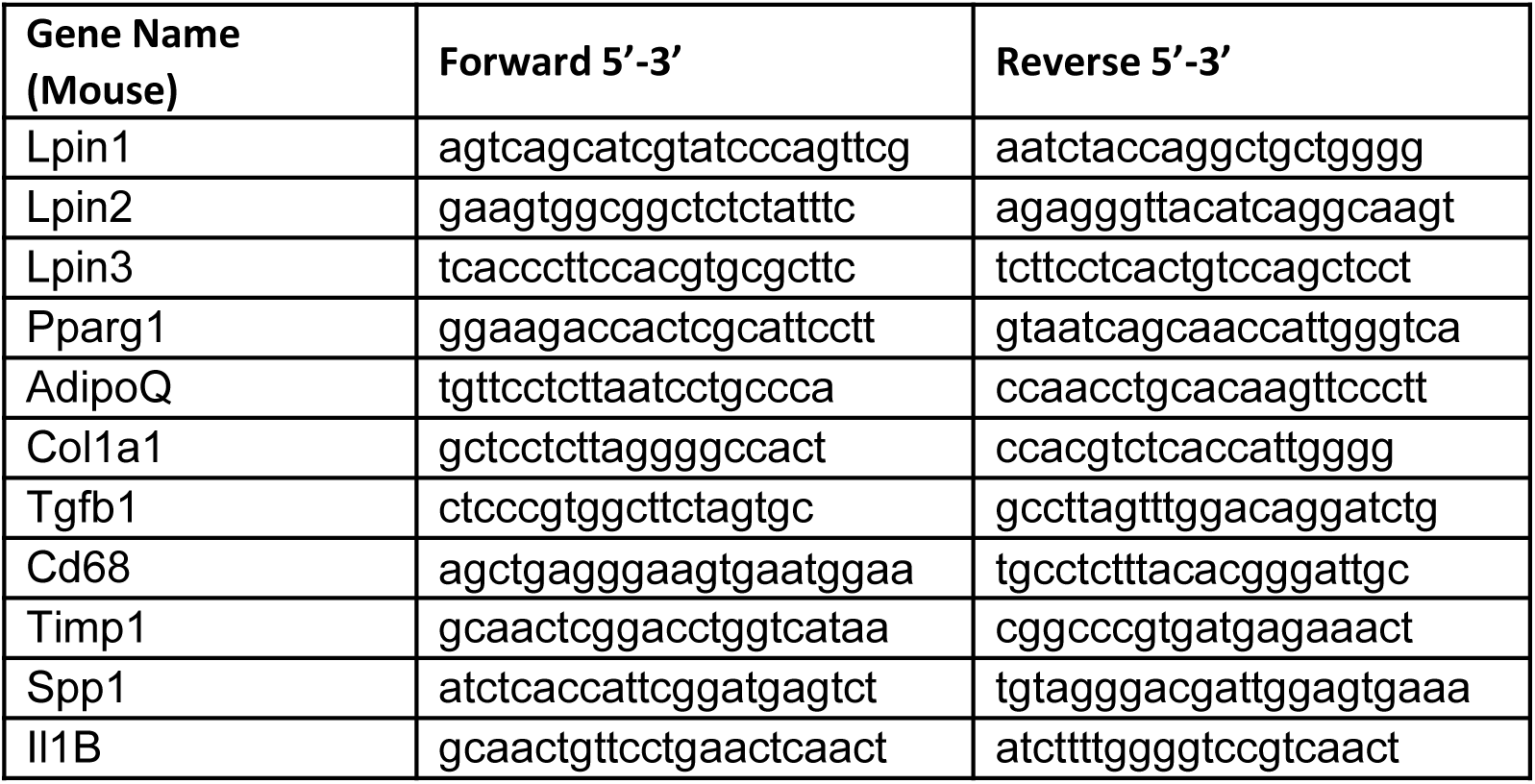
List of primer sequences used for qPCR.

